# Adaptive Integration of Perceptual and Reward Information in an Uncertain World

**DOI:** 10.1101/2024.04.24.590947

**Authors:** Prashanti Ganesh, Radoslaw M. Cichy, Nicolas W. Schuck, Carsten Finke, Rasmus Bruckner

## Abstract

Perceptual uncertainty and salience both impact decision-making, but how these factors precisely impact trial-and-error reinforcement learning is not well understood. Here, we test the hypotheses that (H1) perceptual uncertainty modulates reward-based learning and that (H2) economic decision-making is driven by the value and the salience of sensory information. For this, we combined computational modeling with a perceptual uncertainty-augmented reward-learning task in a human behavioral experiment (*N* = 98). In line with our hypotheses, we found that subjects regulated learning behavior in response to the uncertainty with which they could distinguish choice options based on sensory information (belief state), in addition to the errors they made in predicting outcomes. Moreover, subjects considered a combination of expected values and sensory salience for economic decision-making. Taken together, this shows that perceptual and economic decision-making are closely intertwined and share a common basis for behavior in the real world.

---

In the real world, economic choices fundamentally depend on the processing of perceptual information. An agent first needs to make perceptual decisions, that is, identify the stimuli or states of the environment based on sensory information, to then compute expected values for economic decision-making (Rangel et al., 2008; Summerfield & Tsetsos, 2012). For example, consider a customer who chooses between different types of bread in a bakery. To do so, they need to first identify the available types of bread (states) based on perceptual information to then ascertain the expected taste of the options (expected value). This seemingly simple interplay of perceptual and economic decision-making becomes particularly challenging when perceptual information is ambiguous (perceptual uncertainty) or when outcomes are risky (reward uncertainty) (Bach & Dolan, 2012; Bruckner et al., 2020; Bruckner & Nassar, 2024; Daw, 2014; Ma & Jazayeri, 2014; Platt & Huettel, 2008; Summerfield & Tsetsos, 2012). For example, different loaves of bread might look very similar, yielding perceptual uncertainty. Moreover, the taste of the same type of bread may vary over time or across bakeries due to differences in the ingredients, which leads to reward uncertainty. Therefore, to understand real-world decision-making and learning, we must study the interplay between perceptual and economic choices under uncertainty. Here, we focus on two fundamental questions about this interplay: (i) How does perceptual uncertainty modulate reward learning in humans? (ii) To what extent is human economic decision-making driven by perceptual and value information?

Reward learning requires assigning experienced rewards (e.g., experienced taste after eating a slice of bread) to the states and stimuli of the environment (e.g., type of bread), which is often described as credit assignment (Doya, 2008; O’Reilly & Frank, 2006). This is relatively straightforward when there is clear perceptual information (two distinct types of bread, such as pretzel and baguette; Fig. 1a). However, learning typically takes place amidst perceptual uncertainty due to ambiguous sensory information and internal noise (Walker et al., 2023). In such cases, the state of the environment cannot be clearly identified (Bruckner et al., 2020; Daw, 2014). Therefore, perceptual uncertainty leads to a credit-assignment problem in that the association between reward and state that should be learned is unclear (Fig. 1b). If the decision maker correctly identifies the state, they can accurately learn the association. However, if the decision maker perceives the state incorrectly, they will learn the wrong association between state and outcome.

**Figure 1.**
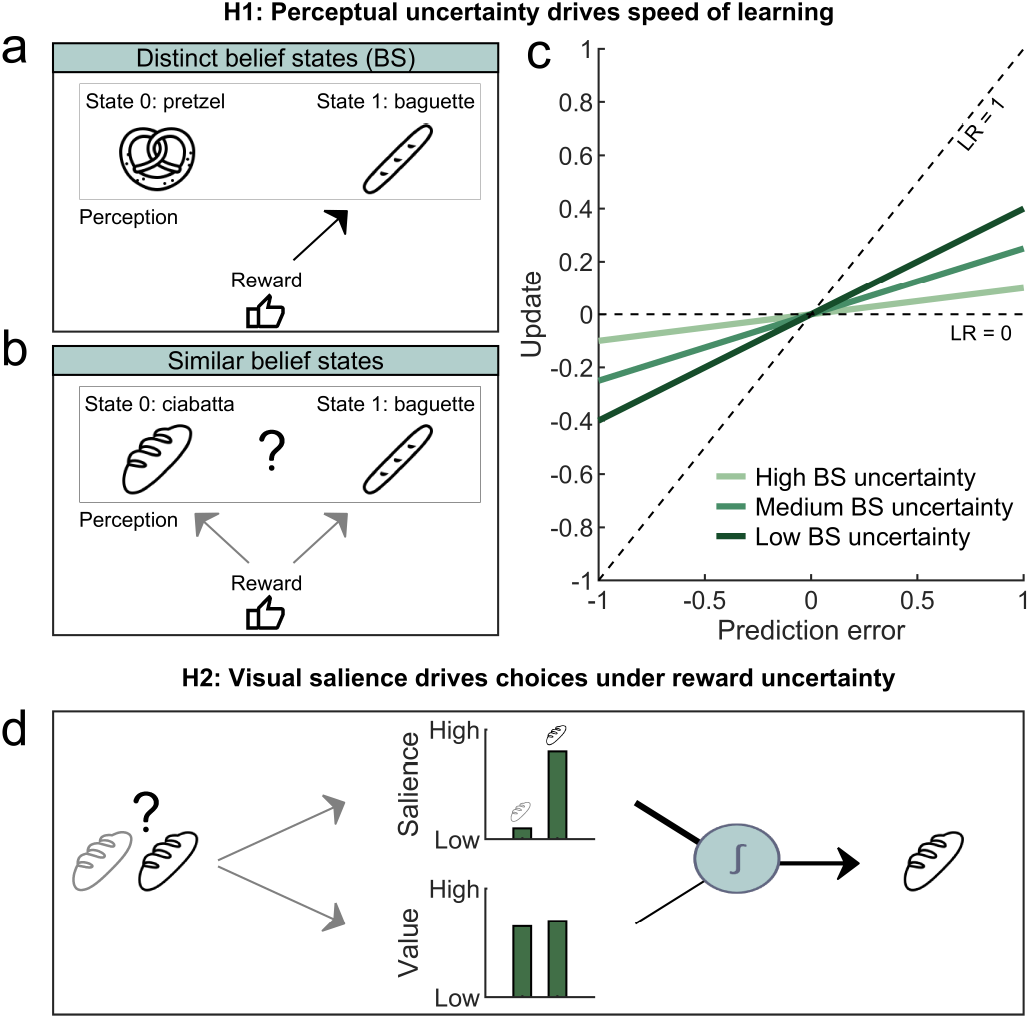
Dynamic integration of visual and reward information under uncertainty. **a**| Learning requires assigning experienced rewards (e.g., taste experience) to the stimuli or states of the environment (e.g., type of bread). In this example, the person can clearly distinguish the two states (pretzel and baguette). When choosing an option (e.g., eating the baguette), they can easily learn an association between reward and state (corresponding to the stimulus “baguette” in this case). **b**| However, when states cannot be clearly dissociated based on sensory information, the person experiences perceptual uncertainty (e.g., two very similar types of bread). In this case, they can compute a belief about the state (belief state), quantifying how confidently the states can be distinguished (e.g., 40% baguette, 60% ciabatta). This leads to a credit-assignment problem, making it unclear what association between state and reward should be updated, and thus, the risk of learning the incorrect association between state and reward. **c**| Our first hypothesis concerns learning under different degrees of uncertainty of belief states. Learning behavior can be quantified using the learning rate (LR; illustrated by the slope of the line). It stands for the rate at which updates about reward expectations change with the prediction error. A learning rate of 1 indicates that only the prediction error is used to make a corresponding update. In contrast, when the learning rate is 0, it indicates that the prediction error has been ignored altogether. We hypothesized that the learning rate tends to be higher, leading to larger updates for a given prediction error when belief states are certain (e.g., 99% baguette, 1% pretzel; dark green line). In contrast, under higher belief-state uncertainty (e.g., 40% baguette, 60% ciabatta; light green line), learning rates are lower. **d**| Our second hypothesis concerns the integration of learned reward expectations (expected value) and visual salience during decision-making. Different options often have, next to different expected values, distinct perceptual features such as salience (e.g., one type of bread captures one’s attention). We hypothesized that both visual salience and expected value govern economic decision-making.

Bayesian inference and reinforcement-learning approaches indicate that the degree of learning from new outcomes should be regulated to deal with perceptual uncertainty (Bruckner et al., 2020; Chrisman, 1992; Ez-zizi et al., 2023; Lak et al., 2017; Larsen et al., 2010). This dynamic regulation of learning behavior is typically quantified by the learning rate. The learning rate expresses to what extent an agent considers the prediction error (i.e., the difference between actual and expected reward) to update their belief about future reward. In doing this, an agent crucially needs to take the probability of being in a particular perceptual state (belief state) into account (Fig. 1a,b). In particular, when the belief state clearly favors a particular state (certain belief state), the learning rate should be higher compared to situations with an uncertain belief state (see light vs. dark green lines in Fig. 1c). In line with these ideas, previous results suggest that humans and animals consider belief states to flexibly regulate learning (Bruckner et al., 2020; Colizoli et al., 2018; Gershman & Uchida, 2019). Moreover, animal work has shown that belief states modulate dopamine activity and choice behavior in perceptual and reward-based decision-making (Babayan et al., 2018; Lak et al., 2017; Lak et al., 2020; Starkweather et al., 2017). However, these findings are primarily based on model fitting of choice data, which does not give direct access to prediction errors, belief updates, and learning rates. Consequently, it only indirectly reveals the impact of belief states on learning. Thus, our goal was to go beyond model fitting by obtaining trial-by-trial measurements of learning and thereby test the direct impact of uncertainty on learning rates (Nassar & Gold, 2013; Nassar et al., 2010; Sato & Kording, 2014). Based on this, we hypothesized that human subjects use lower learning rates when belief states are more uncertain.

The integration of perceptual and reward information is important not only for adaptive learning but also for flexible decision-making under uncertainty. Customers often review bread along different dimensions, such as taste or appearance (artisanal, fluffy), before making a purchase (Fig. 1d). This poses the question of how humans combine perceptual and reward information during economic decision-making. Previous work suggests that humans combine both value information and visual salience of an option to harvest rewards. In many ecological contexts, visual salience elicits species-specific behavior given that they indicate higher levels of safety and certainty (Itti & Koch, 2001; Pike, 2018; Rumbaugh et al., 2007). For instance, a ripe red fruit amidst green leaves reflexively captures one’s attention, thereby increasing the likelihood of survival. Therefore, specifically in perceptually cluttered and uncertain environments, salience could directly modulate economic choices (Navalpakkam et al., 2010; Towal et al., 2013). Based on these considerations, we hypothesized that human economic decision-making is governed by both expected value and perceptual salience.

To test our two hypotheses about the interplay of perception and reward during learning and decision-making, we combined a behavioral choice task with computational modeling. Our results support our first hypothesis that participants adjust their learning rate according to their belief states. In particular, we show that participants use lower learning rates when uncertainty over belief states is higher. This is in line with the predictions of a normative learning model that optimally regulates learning as a function of the belief state. However, next to this normative effect on learning, we also identified a constant effect of prediction errors irrespective of perceptual uncertainty. From the perspective of our model, this effect is sub-optimal, and we interpret it as a heuristic strategy that humans potentially employ to simplify learning. Our results further support our second hypothesis regarding the integration of expected value and salience for decision-making under uncertainty, showing that both drive economic decision-making. Taken together, our study demonstrates how humans integrate perceptual and reward information in the service of adaptive behavior and highlights the convergence of perceptual and economic choices.

## Results

### Task design and performance

To examine the interplay of perception and reward during learning and decision-making, we analyzed the behavioral data of 98 participants (60 male, 38 female; mean age = 23.82 *±* 3.30 standard error of the mean (SEM); range 18-29) completing an online version of the Gabor-Bandit task (Bruckner et al., 2020). Moreover, to optimize the task parameters of the main task, we ran a pilot study with 100 participants (52 female, 48 male; mean age = 22.91*±* 3.04; range 18-30), which we report in the supplement (see Pilot study). Participants were instructed that the goal of the task was to gain as much reward as possible and that each trial comprised three stages (Fig. 2a). In the first stage, participants had to make an economic choice between two Gabor patches. In the second stage, participants received reward feedback on their choice. Finally, in the third stage, they reported their belief about the reward probability using a slider ranging between 0 and 1.

**Figure 2.**
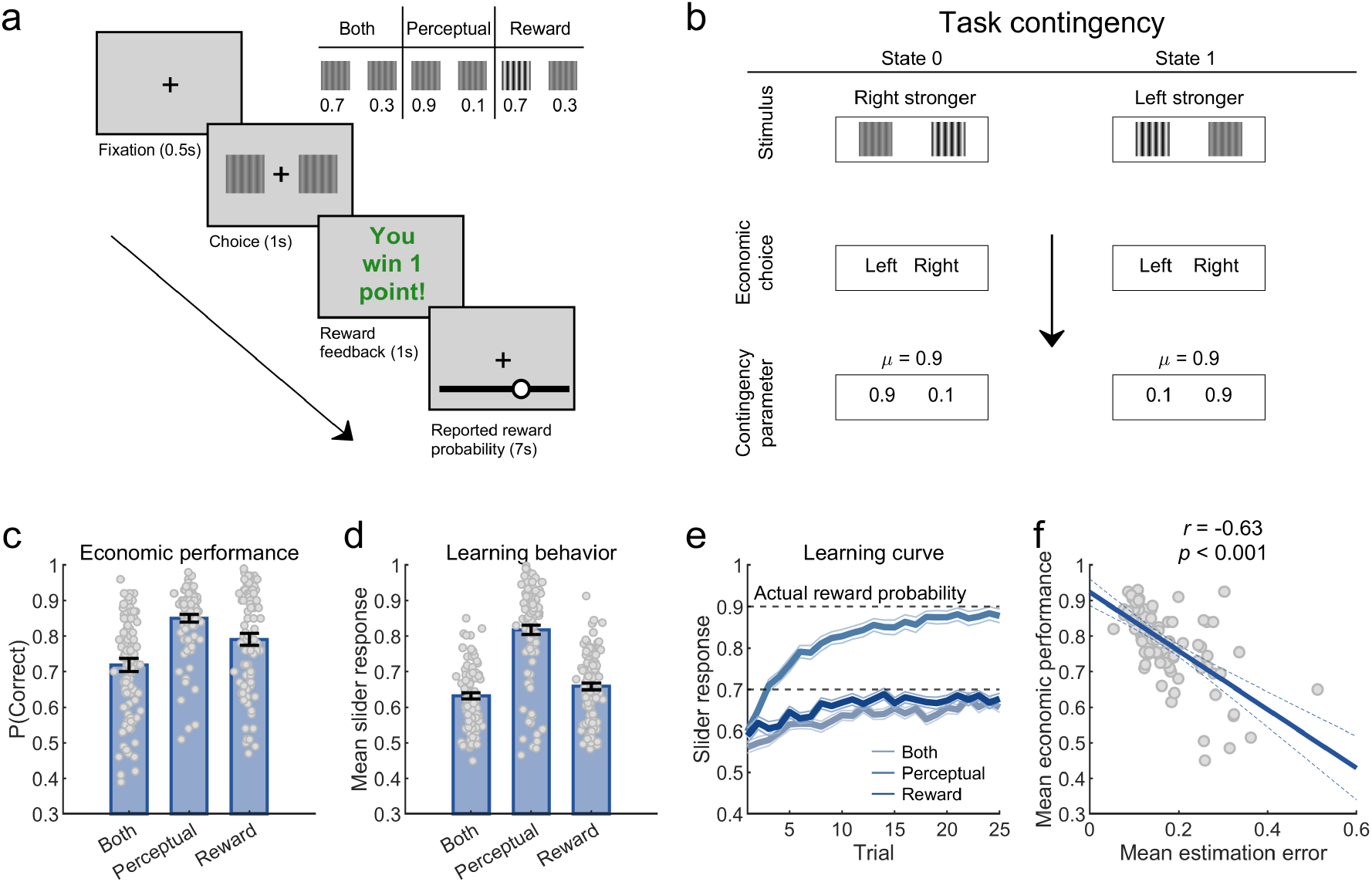
Uncertainty-augmented Gabor-Bandit task, choice performance, and learning behavior. **a**| Subjects were asked to make an economic decision between two Gabor patches. Based on their choice, an outcome was presented. Finally, participants were required to report their subjective value expectation for a hypothetical choice using a slider. **Inset plot**| Experimental conditions. In the “both-uncertainties” condition, participants faced high levels of perceptual uncertainty, where Gabor patches were harder to distinguish, and reward uncertainty, which led to the “correct” option being rewarded with 70 % probability. In the perceptual-uncertainty condition, high levels of perceptual uncertainty were accompanied by low levels of reward uncertainty, which led to the “correct” option being rewarded with 90 % probability. In the reward-uncertainty condition, low levels of perceptual uncertainty, i.e., Gabor patches were easily distinguishable, were combined with high levels of reward uncertainty. **b**| Task contingency. The main aim of the task was to maximize rewards by learning the underlying task contingency between the action and reward, given the state of a trial. Each trial could potentially belong to state 0 or 1. The state determined the location of the high-contrast patch. In state 0, the right patch had a stronger contrast than the left patch and vice versa for state 1. The contingency parameter *µ* determined the reward probability given the action of the participant and the task state. In this example, in state 0, the probability of a reward is higher when choosing the left patch. In state 1, the reward probability is higher when choosing the right patch. Please note that in other blocks, this pattern was reversed, and participants were instructed to relearn the underlying contingency. **c**| Mean *±* standard error of the mean (SEM) economic performance, defined as the frequency of choosing the more rewarding or correct option. **d**| Mean *±* SEM subjective estimate of the reward probability based on the slider responses. **e**| Mean *±* SEM subjective estimate of reward probability based on the slider responses plotted across trials. **f**| Relationship between accuracy in learning (absolute estimation error reflecting absolute difference between true reward probability and slider response) and choice behavior. Lower average estimation errors signal better learning and are moderately correlated with higher levels of economic performance.

Like in classical perceptual decision-making paradigms, the task featured perceptual uncertainty about the Gabor patches (Gold & Stocker, 2017). Moreover, as in classical economic decision-making paradigms, rewards were delivered probabilistically, which is defined as risk or reward uncertainty (Bruckner & Nassar, 2024; Platt & Huettel, 2008; Rangel et al., 2008). On each trial, the patches had varying contrast-difference levels that were determined by a hidden state. In state 0, contrast differences were negative, indicating that the right patch was stronger, while in state 1, contrast differences were positive, and the left patch was stronger. Moreover, the hidden state and reward-contingency parameter governed what economic decision would be rewarded (Fig. 2b). For example, when the contingency parameter assumed the value of 0.9, then in state 0, the left patch with the lower contrast also had a reward probability of 90 %, and the right patch had a reward probability of 10 %. On the other hand, in state 1, the reward contingency was reversed. In this case, the left patch with the higher contrast had a reward probability of 10 % and the right patch of 90 % (see Contingencies, for more details). The participants’ responses on the slider crucially allowed us to track the participants’ beliefs about the reward probability from trial to trial. The task was divided into 12 blocks of 25 trials. The contingency parameter was consistent within each block. Since participants were unaware of the current block’s contingency parameter, they had to learn the parameter value during each block.

To induce perceptual uncertainty and manipulate belief states on a trial-by-trial basis, we manipulated the contrast differences of the patches. The contrast differences were sampled from a uniform distribution. The range of the distributions for high and low perceptual uncertainty was calibrated based on the pilot study. When the contrast differences were small (patches looked more similar), belief states were uncertain. Conversely, for trials in which the two patches had distinct contrast levels, belief-state uncertainty was low. To manipulate reward uncertainty, we manipulated the contingency parameter, where 0.7 (i.e., correct choices rewarded in 70 %) corresponds to higher reward uncertainty and 0.9 (i.e., correct choices rewarded in 90 %) to lower reward uncertainty. The systematic manipulation of uncertainty resulted in three experimental conditions (Fig. 2a inset). The first condition included trials with both forms of uncertainty (termed “both-uncertainties” condition). Consequently, trials with only perceptual or reward uncertainty belong to the perceptual- and reward-uncertainty conditions, respectively. Finally, to ensure that participants had to re-learn the reward contingencies on each block, we counter-balanced the mapping between states, actions, and rewards. That is, in half of the blocks, the patch with the higher contrast level was the rewarding choice option (which we refer to as the high-contrast blocks). In the other half of the blocks, the patch with the lower contrast level was the more rewarding choice option (low-contrast blocks). Please note that this manipulation is crucial since the same mapping between states, actions, and rewards across blocks would negate the need for re-learning after the initial block (see Task details for more details).

To test if participants learned to choose the more rewarding option under both perceptual and reward uncertainty, we analyzed their choices and subjectively reported reward probabilities. Indeed, participants learned to choose the correct option (high-reward option) in all conditions. The average economic choice performance was above chance in all conditions (Fig. 2c; both: mean = 0.72 *±* 0.014, *t*_97_ = 15.79, *p* < 0.001, Cohen’s *d* = 5.23, perceptual: mean = 0.85 *±* 0.009, *t*_97_ = 38.95, *p* < 0.001, Cohen’s *d* = 9.55, reward: mean = 0.79 *±* 0.014, *t*_97_ = 20.1, *p* < 0.001, Cohen’s *d* = 5.52). Moreover, performance was significantly different between the conditions (*F*_2,291_ = 26.77, *p <* 0.001). In line with the intuition that perceptual uncertainty impairs decision-making, economic choice performance in the both-uncertainties condition was lower as compared to the reward-uncertainty condition (*t*_194_ = *−* 3.56, *p <* 0.001, Cohen’s *d* = *−* 0.51). Similarly, the results suggested that reward uncertainty reduced choice performance. Economic choice performance was lower in the both-uncertainties condition than in the perceptual-uncertainty condition (*t*_194_ = *−* 7.92, *p <* 0.001, Cohen’s *d* = *−* 1.13). Choice performance was also better in the perceptual-uncertainty condition as compared to the reward-uncertainty condition (*t*_194_ = 3.51, *p <* 0.001, Cohen’s *d* = 0.5), suggesting that given our experimental settings, the average impact of reward uncertainty was stronger than the impact of perceptual uncertainty on choice performance.

Consistent with the decision-making results, learning curves based on the slider responses clearly demonstrate that participants used the reward feedback to update their beliefs about the reward probabilities (Fig. 2d,e). Participants approached the actual probabilities despite slight underestimation of the probabilities for each condition across trials in a block (both: mean = 0.63*±* 0.01, Cohen’s *d* = 7.26, perceptual: mean = 0.82 *±* 0.01, Cohen’s *d* = 6.12, reward: mean = 0.66 *±* 0.01, Cohen’s *d* = 7.14). There was a significant effect of the type of uncertainty on the mean reported reward probability across the trials in a block (*F*_2,291_ = 87.08, *p <* 0.001). The impact of uncertainty on the reported reward probability was similar to that of its effect on choice behavior. In the both-uncertainties condition, the reported reward probability was lower as compared to the reward-uncertainty condition (*t*_194_ = *−* 2.05, *p* = 0.04, Cohen’s *d* = *−* 0.29), and the perceptual-uncertainty condition (*t*_194_ = *−*11.5, *p <* 0.001, Cohen’s *d* = *−*1.64). Reported reward probability was also higher in the perceptual-uncertainty condition as compared to the reward-uncertainty condition (*t*_194_ = 9.7, *p <* 0.001, Cohen’s *d* = 1.39).

Finally, participants who reported more accurate estimates of the underlying reward probabilities were more likely to make better choices. To quantify the accuracy of participants’ reported reward probabilities, we computed the absolute difference between the actual reward probability and participants’ estimated reward probabilities (estimation error). Lower estimation error indicates higher accuracy of a participant’s belief about the reward probability. Results showed that lower estimation errors were significantly correlated with higher economic choice performance (Fig. 2f; Pearson’s *r*_97_ = −0.63, *p* < 0.001). Building upon these findings about choice and learning behavior we next examined our key hypotheses about the interplay of perceptual and economic choices.

### A normative agent considers belief states to regulate the learning rate

Our first research question is how humans consider perceptual uncertainty during reward-based learning, and we hypothesized that perceptual uncertainty modulates participants’ learning rates. We now illustrate this hypothesis based on simulations using a normative Bayesian agent model that utilizes belief states to regulate learning rates optimally (Bruckner et al., 2020). Akin to human participants, the agent first observes the contrast difference of the Gabor patches. Due to perceptual uncertainty, the agent cannot see the objectively presented difference but instead observes contrasts that are distorted by Gaussian sensory noise (Fig. 3a). Based on its subjective observation, the agent computes the belief state (see Perceptual inference, for more details). Under lower perceptual uncertainty (i.e., less sensory noise), the agent is likely to have more distinct belief states (*π*_*s*_ = (0.1, 0.9)), that is, the agent can identify which patch displays the stronger contrast (Fig. 3b). The normative agent uses the belief states to compute the current expected value of the choice options (see Economic decision-making, for more details). For instance, in the example in Fig. 3c, the learned contingency parameter (*µ* = 1) has a high bearing on the expected values (0.1 for action 0, 0.9 for action 1) since the belief states are highly distinct from one another under lower perceptual uncertainty. In contrast, under higher perceptual uncertainty due to more sensory noise, the agent experiences more uncertain belief states (*π*_*s*_ = (0.4, 0.6)) that lead to discounted expected values (see light green bars in Fig. 3c). Thus, the contingency parameter (*µ* = 1) has lesser influence on expected values under hardly distinguishable belief states, resulting in similar expected values for both actions.

**Figure 3.**
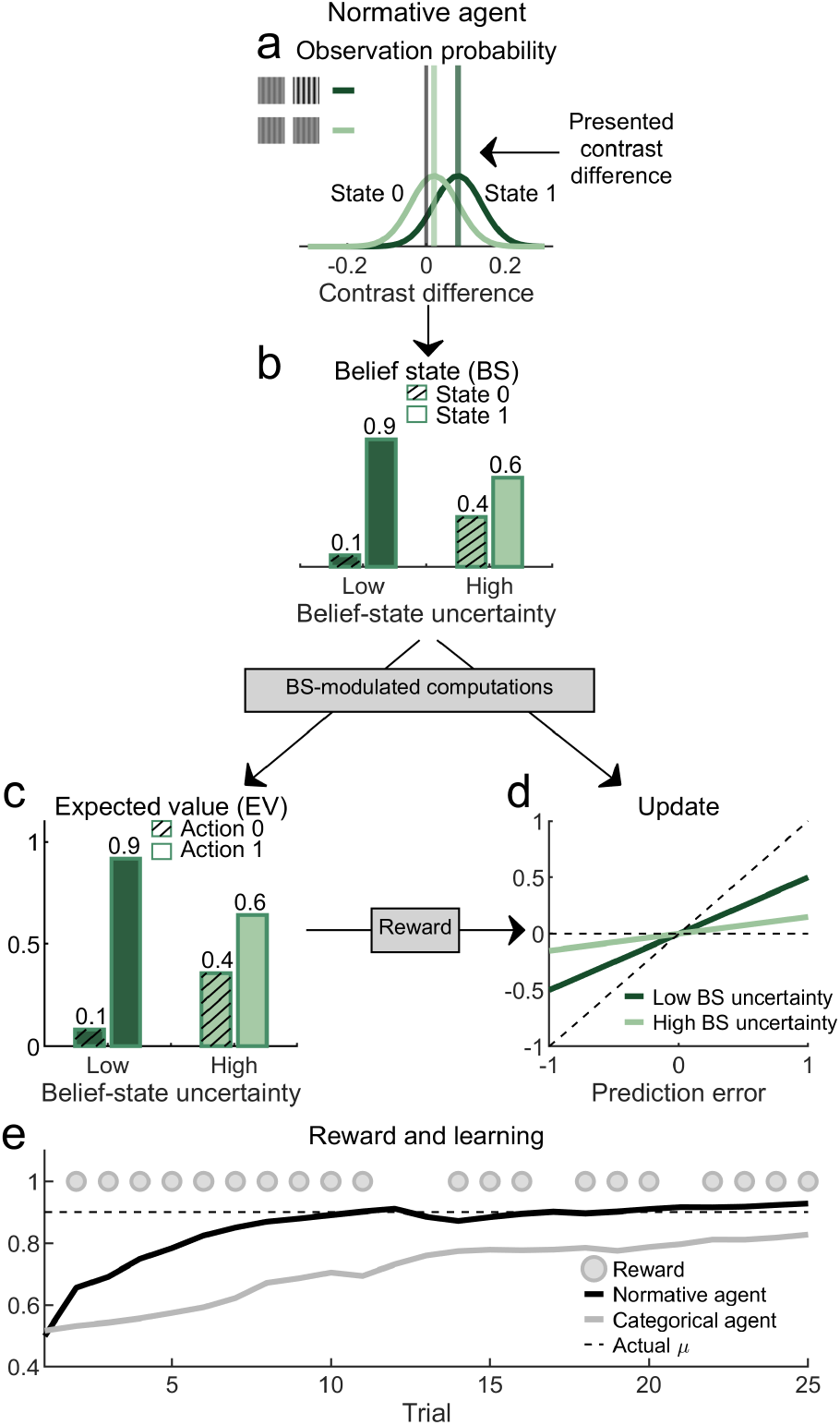
Normative agent. **a**| Contrast-difference observation. A trial can assume one of two hidden task states. The state determines the contrast difference between the high- and low-contrast patches (state *st* = 0 indicates that the right patch has a stronger contrast, and state *st* = 1 indicates that the left patch has a stronger contrast). Due to sensory noise (perceptual uncertainty), the agent cannot perceive the objectively presented contrast difference but a subjective observation that is sampled from a Gaussian observation distribution. Within this distribution, higher perceptual uncertainty is reflected in higher variance over possible observations. **b**| Belief state. The agent computes the probability of being in a given state (belief state) given the subjective observation. Larger contrast differences are translated into more distinct belief states. Subsequently, the agent considers the belief state for economic decision-making and learning. **c**| Uncertainty-weighted expected value. During decision-making, the agent combines the belief state and the learned reward probabilities to compute the expected value. The expected values for the two options are less distinct when belief states are more similar. **d**| Uncertainty-weighted learning. When receiving reward feedback after an economic choice, the agent takes into account the belief state during reward-based learning. The agent uses the belief states to determine how much the prediction error modulates the current trial’s update in the estimate of the contingency parameter. When there is less uncertainty regarding belief states, the agent uses a higher learning rate and, thus, engages in faster learning from prediction errors. However, to deal with the credit-assignment problem arising from highly uncertain belief states (i.e., due to uncertainty, it is unclear what association between stimulus and reward should be updated), the agent dynamically adjusts the learning rate to avoid incorrect assignment of obtained rewards to alternatives. **e**| When learning from multiple outcomes, the estimated contingency parameter approaches the actual contingency parameter with the passage of trials in a block. In contrast, an agent who ignores perceptual uncertainty and represents “categorical” belief states (i.e., assuming that it can perfectly perceive contrast differences and infer the hidden task state) shows reduced learning performance. In this case, the agent often updates the wrong association between stimuli and rewards, thereby leading to an underestimation of the contingency parameter.

Subsequently, the agent makes a choice and receives a reward. Please note that we assumed that the agent’s decisions were free of noise to simulate exploitative choices. We express the underlying learning from the obtained reward as how much the agent updates its belief about the contingency parameter, given the prediction error. When belief states are clearly distinct, the agent uses moderate learning rates (*π*_*s*_ = (0.1, 0.9); see dark green line in Fig. 3d). In contrast, when belief states are more uncertain (*π*_*s*_ = (0.4, 0.6)), the learning rate is considerably lower (see light green line in Fig. 3d). That is, when belief states are ambiguous, the influence of the prediction error on learning from an outcome considerably reduces (see Learning, for more details). Therefore, when perceptual uncertainty is low, the agent makes better choices and learns reward probabilities more quickly (for a comparison between the three conditions, see Fig. S11).

Based on this normative belief-updating mechanism, the agent optimally learns the underlying contingency parameter (see black curve in Fig. 3e). Crucially, considering the belief state in this way during learning yields a more accurate belief about the contingency parameter compared to a learning mechanism that ignores perceptual uncertainty. Specifically, the learning curve of an agent that only represents binary or categorical belief states (belief states only assume 0 and 1 instead of values in between) reflects a less accurate and biased belief about the contingency parameter (categorical agent; see gray curve in Fig. 3e). In summary, these simulations illustrate our first hypothesis that the certainty of a belief state modulates learning rates. When belief states are more certain, learning rates tend to be higher than on trials with more uncertain belief states.

### Humans consider belief states to regulate the learning rate

We next tested our first hypothesis that humans take into account their belief states to regulate learning behavior. We quantified participants’ learning behavior on each trial by calculating the learning rate. To do so, we used the reported beliefs about the reward probability to compute each trial’s prediction error and belief update (see Data preprocessing, for more details). Sub-sequently, learning was measured as the extent to which participants updated their subjective estimate of the reward probability on the slider, given that trial’s prediction error. We approx-imated belief states using the level of contrast difference, where lower differences result in more uncertain belief states.

Directly comparing single-trial learning rates across bins of contrast-difference values (ordered from more to less uncertain approximated belief states), we observed an increase in the learning rate (Fig. 4a). That is, participants learned more when belief states were, on average, more certain, in line with our hypothesis that perceptual uncertainty leads to dynamic adjustment of learning rates.

**Figure 4.**
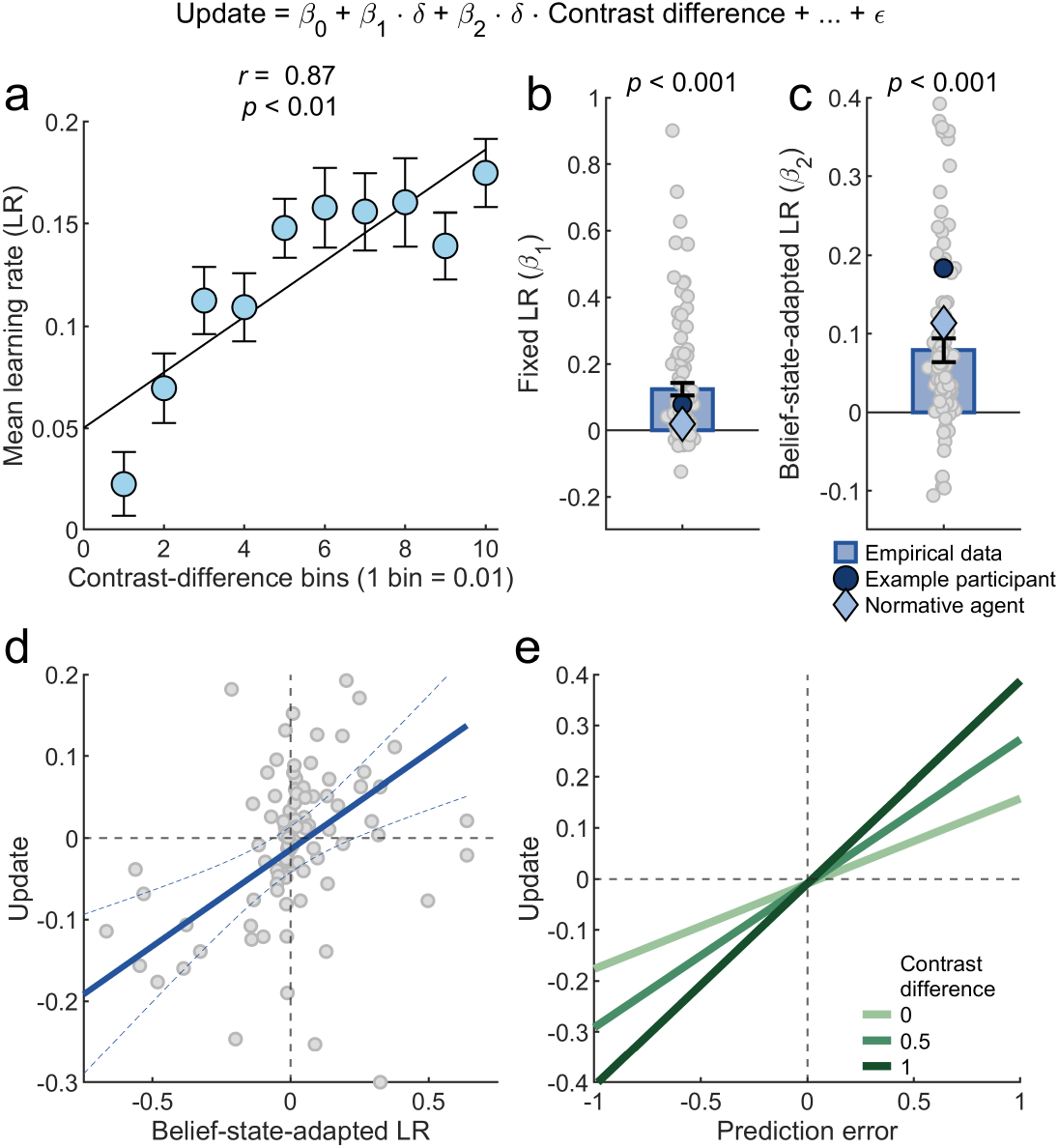
Decomposing learning rates. **a**| We computed single-trial learning rates reflecting the extent to which prediction errors (difference between obtained reward and subjectively reported reward probability) drive slider updates (difference in reported reward probability between current and previous trial). To examine whether learning rates were dynamically adjusted to how well subjects could discriminate the choice options, we divided the data into 10 contrast-difference bins, where lower bins correspond to more uncertain belief states. The plot shows the mean*±* standard error of the mean (SEM) learning rate for each bin. The increase as a function of contrast difference (Pearson’s *r*_08_ = 0.87, *p* = 0.001) suggests that subjects use higher learning rates when belief states are more clearly distinct. **b**| To decompose the influences of different factors on the learning rate, we developed a regression model (see inset equation on top of plot, where *δ* denotes the prediction error). Mean *±* SEM coefficients for key regressors from the linear regression model are shown here. Positive fixed-LR coefficients indicate participants’ average tendency to learn from prediction errors (Cohen’s *d* = 0.68). **c**| The belief-state-adapted-LR coefficients reflect the adjustment of the learning conditional of the contrast difference (Cohen’s *d* = 0.53). **d**| This subplot shows an example participant illustrating the extent to which prediction errors weighted by contrast difference (belief-state-adapted LR) drive the update. In line with (a), this suggests a down-regulation of the learning rate when belief states are more uncertain. **e**| Across three levels of contrast-difference values, regression fits for a range of prediction errors of an example participant suggest that belief states modulated the learning rate. Higher contrast differences (i.e., on average, more distinct belief states) led to larger updates as compared to lower contrast differences.

While the previous analysis suggests uncertainty-driven belief updating on the group level, it does not indicate to what extent individual subjects use the belief state to weight the prediction error. Therefore, we used a linear regression model that quantified the impact of prediction errors and belief states on belief updating for each subject (McGuire et al., 2014; Nassar et al., 2019; Sato & Kording, 2014). In the model, we expressed the reported belief update as a linear function of the prediction error, and the slope of this function is equivalent to a fixed learning rate as in typical error-driven learning models (often referred to as *α* in reinforcement learning; Daw, 2014). To model the dynamic impact of belief states, the model included an interaction term between belief state and prediction error (Fig. 4 inset equation, where *δ* denotes prediction error). The model also allowed us to simultaneously control for the impact of choice confirmation and several nuisance variables (for more details on the model, see Regression analysis). We fit the model to participants’ single-trial updates as well as simulated data based on the normative agent. Comparison with the predictions of the model allowed us to ascertain to what extent human learning under uncertainty approaches normative belief updating.

Participants’ fixed learning rates reflecting the average influence of prediction errors were positive (mean = 0.12*±* 0.018, *t*_97_ = 6.72, *p <* 0.001, Cohen’s *d* = 0.68; Fig. 4c, fixed-learning-rate (LR) coefficient). A systematic comparison to the normative agent suggests that participants’ positive fixed learning rates correspond to a heuristic, if not a biasing influence of the prediction error on learning. The agent shows a coefficient near zero, indicating that from a normative learning perspective, learning behavior should not be driven by a static influence of prediction error, thus leaving room for uncertainty-driven flexible learning.

Besides the overall effect of the prediction error, participants showed evidence of dynamic learning-rate adjustments similar to the normative model. We found that larger contrast differences (i.e., on average, more certain belief states) propelled updates for a given prediction error, as indicated by the positive coefficients for the interaction of prediction error and contrast difference (mean = 0.08 *±* 0.015, *t*_97_ = 5.23, *p <* 0.001, Cohen’s *d* = 0.53; Fig. 4d, belief-state-adapted-LR coefficient). That is, in accordance with the normative model, participants flexibly adjust their learning rate depending on the belief state. Despite considerable hetero-geneity across participants, on average, participants align with the agent’s prescriptions to take perceptual uncertainty into account.

Follow-up analyses of the belief-state-adapted-LR coefficient suggested a small but concise influence on the learning rate. Expressing the relationship between this coefficient and the updates, after taking all other regressors into account, a robust relationship between the belief-state-weighted prediction errors and updates is evident. To illustrate this point, Fig. 4d shows an example participant whose regression coefficient is indicated by the blue dot in Fig. 4c. This plot shows that for a positive coefficient, the belief update systematically increases with contrast difference. Furthermore, we plotted the relationship between prediction errors and updates for varying contrast-difference levels for the same example participant (Fig. 4e). This analysis similarly shows that for a given prediction error, learning rates systematically increase with decreasing belief-state uncertainty (increasing contrast-difference values). Please refer to Regression diagnostics in the methods for more details and for additional information on other regressors, see Full learning-rate analysis. Moreover, we found evidence for a preference to learn more strongly from outcomes that confirm choices, suggesting the presence of a choice-confirmation bias. In our regression model, positive choice-confirmation coefficients indicate stronger updates following prediction errors computed after receiving reward feedback that confirms choices (mean = 0.07 *±* 0.009, *t*_97_ = 8.03, *p <* 0.001, Cohen’s *d* = 0.81, Fig. S2a, confirmation bias). Finally, in our current approach, the coefficients for the belief-state-driven learning could either be due to (i) a strategic calibration of the learning rate to perceptual uncertainty or (ii) state confusion due to perceptual uncertainty. To tease these apart and focus on update magnitude, we fit the same model to absolute updates with absolute prediction errors (for more details, see Absolute learning-rate analysis). Our results from this approach were consistent with the aforementioned results (Fig. S1c). In conclusion, our combined analyses suggest that reward learning under perceptual uncertainty is molded by the belief state.

### Fixed but not flexible learning impacts belief accuracy

Thus far, our results suggest that humans adaptively adjust their learning rates under perceptual uncertainty. However, are individual differences in the degree of learning flexibly associated with the accuracy of a participant’s beliefs? An obvious benefit of such belief-state-adapted learning is that beliefs are less likely to be corrupted by perceptual uncertainty. One crucial question that follows from this is whether individual differences in the degree of flexible learning translate into differences in the accuracy of beliefs. To investigate this, we employed an exploratory approach to predicting average estimation error (absolute difference between the actual reward probability and the value reported by the participant) based on the fixed and flexible learning-rate coefficients from our regression analysis.

We found that subjects with high fixed learning-rate coefficients (i.e., prediction-error-driven learning) tended to have larger estimation errors (*β* = 0.65, *p <* 0.001; Fig. 5a). In a stable environment, such as in our task, rash learning has adverse effects on belief updates as it is linked to large shifts in estimates and, possibly, stronger deviations from the actual reward probability. In contrast, subjects who made smaller learning adjustments (indicated by low and moderate fixed-LR coefficients) consequently reported more accurate estimates. However, individual differences in belief-state-adapted-LR coefficients did not have a significant relationship with estimation error (*β* = 0.09, *p* = 0.37; Fig. 5b). One explanation for the absence of an effect of the belief-state-adapted LR might be the strong biasing effect of the fixed LR on estimation errors that could potentially overwrite its influence. However, we found that absolute belief-state-adapted LRs have a significant relationship with belief accuracy. One key reason for this could be that absolute learning rates better capture strategic calibration of learning under uncertainty, hence are linked to more accurate beliefs (see Absolute learning and estimation error and Fig. S9b). Similarly, we found no significant links between estimation error and confirmation bias (*β* = 0.19, *p* = 0.055; Fig. 5c). We also found that the confirmation bias is linked with more accurate subjective estimates of the reward probability (see Confirmation bias and over-estimated beliefs and Fig. S7). See Signed learning rate and estimation error for details on other signed learning regressors.

**Figure 5.**
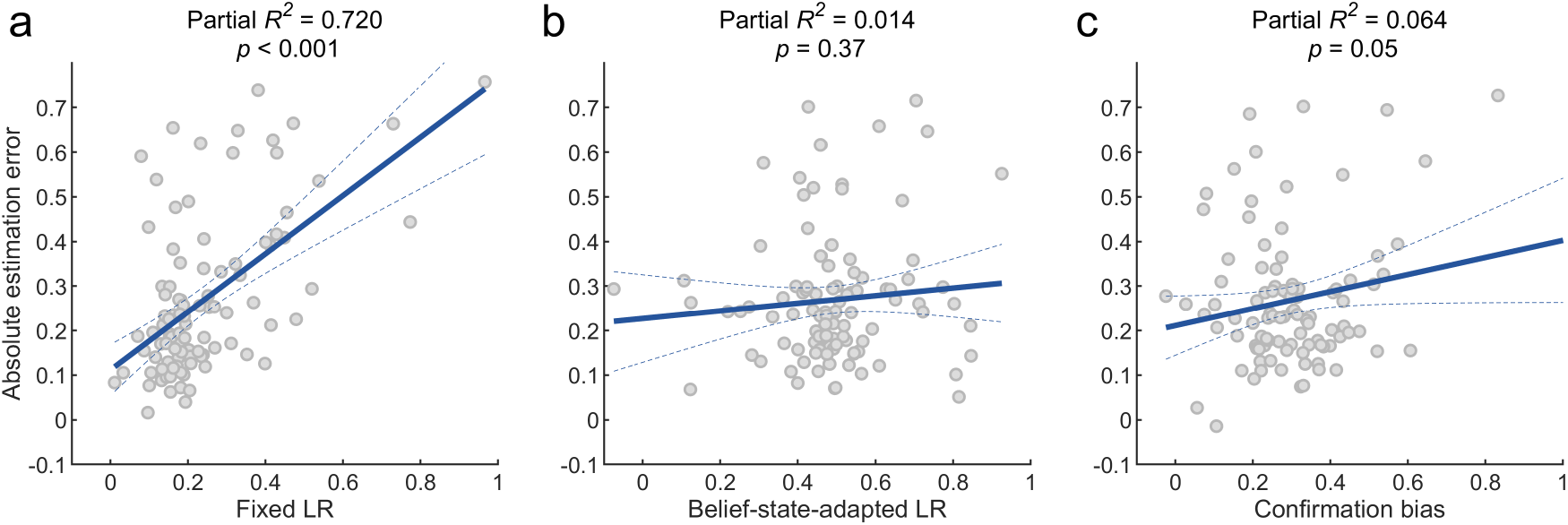
Influence of learning on belief accuracy. We examined the relationship between absolute estimation errors (difference between actual reward probability and subjective estimate of the probability) reflecting belief accuracy and several predictors of the regression model (fixed learning rates, belief states, confirmation bias). **a**| Larger fixed learning rates were associated with larger absolute estimation errors, suggesting that learning too much from a prediction error negatively impacts learning. We did not find a systematic effect of **b**| belief-state-adapted learning rates and **c**| the confirmation bias.

### Economic choices are governed by expected values and visual salience

We next tested our second hypothesis that both value and visual salience govern economic decision-making. In the context of our task, we assume that options with higher contrast levels have a higher perceptual salience than the options with lower contrast. To quantify a hypothetical effect of salience on economic decision-making, we compared economic choice performance between high- and low-contrast blocks. Please recall that in half of the blocks of our task, the high-contrast option yielded more rewards (high-contrast blocks), and in the other half, the low-contrast option was more rewarding (low-contrast blocks). Therefore, a higher economic choice performance in high-contrast than low-contrast blocks reveals a “salience” bias towards the more salient option, indicating a combined impact of perceptual and reward information as hypothesized. In contrast, an alternate hypothesis would state the absence of this bias, which translates to a similar reward-maximizing performance for high- and low-contrast blocks. This analysis indicated that participants showed a significant salience bias in the both-uncertainties (mean = 0.06 *±* 0.025, *t*_97_ = 2.61, *p* = 0.01, Cohen’s *d* = 0.26) and reward condition (mean = 0.1*±* 0.02, *t*_97_ = 5.02, *p <* 0.001, Cohen’s *d* = 0.51). However, as hypothesized, in the perceptual-uncertainty condition (mean = 0.02*±* 0.012, *t*_97_ = 1.8, *p* = 0.08, Cohen’s *d* = 0.18), we did not find a significant salience bias (Fig. 6a).

**Figure 6.**
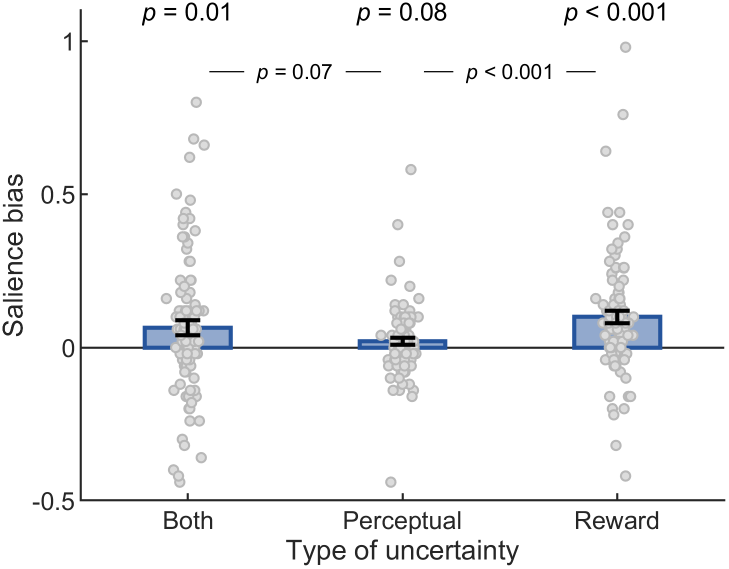
Salience bias. We examined whether choice performance was governed by perceptual salience. In high-contrast blocks, the more salient option had a higher reward probability and vice versa for low-contrast blocks. Therefore, higher choice performance on high-contrast than low-contrast blocks reflects a positive choice bias towards the more salient option. The plot shows the mean *±* standard error of the mean (SEM) salience bias (difference in economic performance between high- and low-contrast blocks) for the different types of uncertainty, which is significant in the condition with both perceptual and reward uncertainty (both-uncertainties condition) and the reward-uncertainty condition but not in the perceptual-uncertainty condition.

Next, we examined if the salience bias is more enhanced due to reward uncertainty by comparing the salience bias in high reward uncertainty (“both” and reward uncertainty) blocks with low reward uncertainty (perceptual uncertainty) blocks. Participants showed a significantly larger salience bias in the reward-uncertainty condition as compared to the perceptual-uncertainty condition (*t*_97_ = 4.04, *p <* 0.001, Cohen’s *d* = *−* 0.49). However, participants did not show a significantly pronounced salience bias in the both-uncertainties condition as compared to the perceptual-uncertainty condition (*t*_97_ = 1.82, *p* = 0.07, Cohen’s *d* = *−*0.23). Pilot study results also showed a salience bias which is modulated by the extent of reward uncertainty (Fig. S10). Overall, these findings suggest that participants’ decisions are driven by both expected values and visual salience, and we identified reward uncertainty as a facilitating factor for the same.

## Discussion

In an uncertain world, the interplay of perceptual and reward information is crucial for adaptive behavior. To study this, we introduced an uncertainty-augmented task combining perceptual and economic decision-making that allows for the direct estimation of the learning rate in a trial-by-trial fashion. Combined with computational modeling, we found that uncertainty plays a key role in integrating perceptual and economic decision-making. First, we show that humans flexibly modulate learning rates according to the uncertainty over the distinguishability of choices based on sensory information (belief state). Thus, this provides crucial evidence for our first hypothesis (H1) that perceptual uncertainty drives the speed of reward learning. Second, we found that humans show a choice bias towards the more perceptually salient option. This aligns with our second hypothesis (H2) that reward uncertainty facilitates the combined impact of perceptual and reward information on choices. Together, our results emphasize the intertwined nature of perceptual and economic decision-making.

As hypothesized, we showed that humans adjust the learning rate in response to varying belief states. When sensory information was more ambiguous, and belief states were presumably more uncertain, subjects updated their estimates of expected values to a lesser extent, in line with a reduced learning rate. Under perceptual uncertainty, identifying stimuli and environmental states is difficult (Bach & Dolan, 2012; Ma & Jazayeri, 2014; Rao, 2010), which makes it challenging to assign experienced rewards to the correct state during credit assignment (Babayan et al., 2018; Courville et al., 2006; Doya, 2008; O’Reilly & Frank, 2006; Rao, 2010). Our results suggest that to avoid incorrect pairing of state and reward, humans resort to a belief-state-guided learning strategy (Bruckner et al., 2020; Chrisman, 1992; Ez-zizi et al., 2023; Lak et al., 2017; Lak et al., 2020; Larsen et al., 2010; Starkweather et al., 2017).

Our results on learning-rate adjustments to perceptual uncertainty go beyond the domain of perceptual estimation and show that this mechanism is transferable to reward learning. Sato and Kording (2014) and Vilares et al. (2012) used a continuous perceptual estimation task in which visual targets had to be predicted based on uncertain sensory information. Subjects adjusted predictions to a lesser extent when perceptual uncertainty was higher, which aligns with our results despite key differences. Crucially, in these studies, perceptual uncertainty originated from external noise inherent to the presented information, while in our task, perceptual uncertainty is primarily due to the imprecision in the human visual system (exact contrast differences are hard to detect for humans). Moreover, in our work, subjects had to learn reward probabilities under perceptual uncertainty from binary rewards, as opposed to perceptual estimation. Together, these lines of research converge on the view that this mechanism is a ubiquitous phenomenon that generalizes across different scenarios.

Moreover, the findings from Drevet et al. (2022) are in line with our results regarding the regulation of learning rates to stimulus discriminability. However, the suggested mapping between belief state and learning rate differs between the studies. Most importantly, Drevet et al. (2022) found evidence of a belief-state threshold above which perceptual information is deemed to be strong and certain enough for learning. Below this threshold, newly arriving information is discarded, which differs from our more continuous down-regulation of learning in response to the belief state. However, there are crucial methodological differences. While Drevet et al. (2022) exposed participants to a changing environment and used binary-choice data to estimate learning dynamics based on model fitting, we used direct reports of belief updating in a stable environment. Future work could combine our direct learning-rate measurements and the extensive model space of Drevet et al. (2022) to compare the two explanations in a common study.

However, our results appear to be at odds with work suggesting that belief-state-driven flexible learning does not occur under perceptual uncertainty in a volatile environment (Ez-zizi et al., 2023). This could be explained by at least two reasons. One key difference to our study is how perceptual uncertainty was induced. Whereas in our work and other previous studies (Bruckner et al., 2020; Lak et al., 2017; Lak et al., 2020; Sato & Kording, 2014; Vilares et al., 2012), perceptual information was associated with varying degrees of belief-state uncertainty, participants in Ez-zizi et al. (2023) were presented with fixed stimuli calibrated to a pre-defined accuracy level. This potentially leaves little room and need for fine-tuning of learning. Moreover, the computational model did not explicitly assume that reward probabilities changed throughout the task, potentially resulting in a worse model fit (Ez-zizi et al., 2023; Larsen et al., 2010). Future work could explicitly incorporate environmental changes into these models to further investigate the interplay of perceptual uncertainty and surprise (Bruckner et al., 2022).

A relevant topic for future research based on our findings is examining the psychophysiological mechanisms behind uncertainty-led flexible learning. Different forms of uncertainty have been linked to the arousal system (Aston-Jones & Cohen, 2005; Yu & Dayan, 2005). In particular, studies using pupillometry as a proxy of arousal suggest that arousal modulates the influence of incoming information on learning (Nassar et al., 2012), perceptual (Krishnamurthy et al., 2017), and choice (de Gee et al., 2017; Urai et al., 2017) biases. One potential neural mechanism behind these effects is the locus coeruleus-norepinephrine (LC-NE) system (Aston-Jones & Cohen, 2005; Gilzenrat et al., 2010; Joshi et al., 2016; Megemont et al., 2022; Murphy et al., 2014; Murphy et al., 2011; Reimer et al., 2016). Therefore, future work could examine the link between arousal dynamics, learning-rate adjustments, and perceptual uncertainty.

Another avenue for future work is improving the slider design that we used to measure learning. We present analyses examining the split-half reliability of our parameters (see Split-half reliability for more details). We found moderately correlated fixed learning rates but weaker correlations for flexible learning (Fig. S6). These values seem to be comparable to similar state-of-the-art Bayesian and reinforcement-leaning approaches and sufficient for group-level analyses of healthy subjects (Loosen et al., 2022; Palminteri & Chevallier, 2018; Patzelt et al., 2018; Schaaf et al., 2023). However, applying our approach to clinical populations or studies interested in individual differences would particularly benefit from more stable estimates. Among many factors that impact reliability, Schurr et al. (2024) identify random noise in behavioral measurements that could arise from discrepancies given the current application of the slider. Improvements to the slider design, including cues about previous estimates (to reduce motor noise) and modifying the starting point of the slider (requiring fewer adjustments), could increase the overall reliability of the parameters.

Furthermore, our second aim was to examine how economic decision-making is swayed by perceptual and reward information. We hypothesized that visual salience, next to the established role of expected values (Bartra et al., 2013; Kable & Glimcher, 2009; Levy & Glimcher, 2012; Rangel et al., 2008), impacts choices. Confirming this, we identified a salience bias that led to a preference for choice options with stronger as opposed to weaker contrasts. These findings suggest that people use salience as a proxy for expected value when value information is uncertain. This result aligns with studies reporting effects of both value and salience in a perceptual choice task (Navalpakkam et al., 2010; Towal et al., 2013). More generally, salience does impact decisions that should ideally be driven solely by value because of its evolutionary significance (Itti & Koch, 2001; Pike, 2018; Rumbaugh et al., 2007) and hence, may be deemed more rewarding in risky environments.

To summarize, we found that humans effectively integrate uncertain perceptual and reward information for learning and decision-making. Humans dynamically adjust reward learning contingent on perceptual uncertainty. Moreover, perceptual salience, in addition to the expected value, drives economic decision-making, where the interaction is guided by reward uncertainty. These findings offer insight into mechanisms behind the interplay of perceptual and reward information, highlighting that each is not solely tied to either perceptual or economic decision-making.

## Methods

### Participants

The study included two experiments. 100 participants were recruited for the main task (38 female, 62 male; mean age = 23.82 *±* 3.30 SEM; range 18-29). All participants were recruited via Prolific (www.prolific.co) for online behavioral experiments. Participants provided informed consent before starting the experiments. We applied several inclusion criteria using Prolific’s participant pre-screen tool. Participants had to be between the ages of 18 and 30 and have normal or corrected-to-normal vision. Additionally, only participants who reported not having used any medication to treat symptoms of depression, anxiety, or low mood were recruited. We did not recruit participants reporting mild cognitive impairment, dementia, and autism spectrum disorder. For taking part in the study, participants were paid a standard rate of £6.00. Moreover, to incentivize their performance, participants received an extra bonus payment of up to £2.50, determined by their economic choice performance. The study was approved by the ethics committee of the Department of Education and Psychology at Freie Universität Berlin (“Effects of Perceptual Uncertainty on Value-Based Decision Making”, protocol number: 121/2016). Data from two participants was rejected since they performed with less than 50% accuracy.

### Experimental task

### Experimental procedure

The task was programmed in JavaScript using jsPsych (version 6.3.0). The Gabor-Bandit (GB) task version of this study comprised three stages (economic decision-making, reward feedback, slider response) (Fig. 2a). In the first stage, the stimulus material comprised a fixation cross and two sinusoidal gratings presented on a screen with a gray background color (#808080). These were created using HTML canvas, which is an in-built element of JavaScript that allows for dynamic rendering of 2-dimensional graphics. To create the gratings, we use a sine texture consisting of two alternative bands of black (#000) and white (#FFF) color with a spatial frequency of 2 cycles per cm. The orientation of the patches was kept constant at 0°.

To manipulate the Gabor-patch contrasts *g*, we controlled the patches’ visibility *v*, where 0 indicates that the patch is transparent (equal to the background) and 1 that it is fully opaque. Subsequently, the displayed contrast of each patch was a weighted combination of the stimulus properties *z* (as defined by the HTML canvas settings described above) and the background color *h*: *g* = *vz* + (1*− v*)*h*. The mean visibility of both patches was maintained at *v* = 0.5. The choice gratings were presented for 1000 ms, and the fixation cross remained on the screen throughout. During the stimulus presentation, participants were required to make the economic choice using the left and right cursor buttons of the computer keyboard. The participants’ responses did not end the patch presentation. In the second trial stage, participants were presented with feedback of winning either zero (“You win 0 points!”) or one (“You win 1 point!”) point based on their economic choice for 1000 ms. Finally, it included an additional probe phase during which participants reported their subjective estimate of the reward probability for a hypothetical choice using a slider. Participants completed 25 trials in each block of the task. The presentation order of blocks was randomized across participants. If a participant failed to respond to a trial, the same trial was repeated at the end of the block.

### Task contingencies

The central feature of our task is that each block of trials inherently features a particular stateaction-reward association. A trial could potentially belong to one of the two hidden task states *s*_*t*_ *∈*{0, 1}. When a trial belongs to *s*_*t*_ = 0, the patch on the left side of the fixation cross has a lower contrast level than the right patch. This relationship is reversed when the trial belongs to *s*_*t*_ = 1. Half of the trials in one block belonged to *s*_*t*_ = 0, while the other half belonged to *s*_*t*_ = 1. Moreover, we refer to the two choices (left vs. right patch) as actions *a*_*t*_ *∈*{0, 1}, where *a*_*t*_ = 0 indicates choosing the left patch and *a*_*t*_ = 1 the right patch. The reward probabilities depended on the state-action combination. For example, when the left patch had the lower contrast level (*s*_*t*_ = 0) and was chosen by the participant (*a*_*t*_ = 0), it was more likely that the participant would obtain a reward. Similarly, when the right patch had the lower contrast (*s*_*t*_ = 1) and was chosen by the participant (*a*_*t*_ = 1), it was likely to yield a reward. In contrast, when in state *s*_*t*_ = 0 (left patch has the lower contrast) and choosing action *a*_*t*_ = 1 (right patch) or when in state *s*_*t*_ = 1 (right patch has the lower contrast) and choosing *a*_*t*_ = 0 (left patch), the reward probability is low. Thus, in such blocks, the low-contrast patch was the more rewarding option (low-contrast blocks). Importantly, in half of the blocks, the state-action-reward contingency was reversed i.e., the high-contrast patch was the more rewarding option (high-contrast blocks). The block order was randomized, and hence, the reward contingencies had to be relearned on each block. Consequently, participants were required to learn the correct association between Gabor-patch locations (states), choices (actions), and obtained rewards to maximize their outcome.

### Task details

The main task comprised 12 blocks with 25 trials and featured three conditions: the “both-uncertainties” condition, the perceptual-uncertainty condition, and the reward-uncertainty condition. In the “both-uncertainties” and perceptual-uncertainty conditions, the contrast difference of two patches was randomly sampled from a uniform distribution of [−0.1 to 0] when the trial belonged to *s*_*t*_ = 0 with the left patch having the lower contrast and [0 to 0.1] when the trial belonged to *s*_*t*_ = 1 and the right patch had the lower contrast. Thus, the absolute contrast levels of the patches ranged from 0.40 to 0.60. In the reward-uncertainty condition with low perceptual uncertainty, the contrast difference of the two patches was in the range −0.35 to −0.45 for *s*_*t*_ = 0 and 0.35 to 0.45 for *s*_*t*_ = 1. Thus, the absolute range of contrast levels was 0.05 to 0.95.

Crucially, in the slider probe phase, the patches were clearly distinguishable; that is, participants did not experience uncertainty about the task state. To report their estimates of the reward probabilities, participants were instructed to click and drag across the slider that ranged from 0 to 100%. To ensure that exclusive use of the more rewarding option as the hypothetical choice does not help participants to learn the state-action-reward contingency, we ask them to report the estimated reward probabilities for both the more and less rewarding option across blocks in the task. In half of the blocks, the hypothetical choice was congruent with the more rewarding patch in the given block (congruent blocks). That is, in half of the high-contrast blocks, the hypothetical choice during the slider phase was congruent with the more rewarding option. Thus, the participants were asked to report their subjective estimate for the high-contrast patch (more rewarding option). However, on the other half of the high-contrast blocks, participants were asked to report for the low-contrast option (less rewarding option). That is, the hypothetical choice was incongruent with the more rewarding patch in that block of trials (incongruent blocks). Finally, the order of task blocks was randomized for each participant.

### Gabor-Bandit task model

We simulated predictions of a Bayes-optimal learning model (Bruckner et al., 2020). To describe the model in detail, we first present a model of the Gabor-Bandit task. In line with Bruckner et al. (2020),

- *T* := 25 indicates the number of trials per block, where we use *t* as the trial index,
- *S ∈*{0, 1} denotes the set of task states, where 0 indicates that the right patch has stronger contrast than the right patch and vice versa for state 1; the state also determines the action-reward contingency in the task,
- *C∈* [*− κ, κ*] is the set of contrast differences between the patches, where *κ* indicates the maximal contrast difference, which differs across conditions in this work, as described above,
- *A∈*{0, 1}refers to the set of economic choices, where 0 refers to choosing the left patch, and 1 refers to choosing the right patch,
- *R ∈* {*0, 1*} denotes the set of rewards,
- *p*^*ϕ*^(*s*_*t*_) is the Bernoulli state distribution defined by

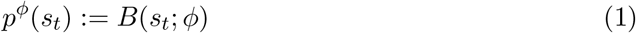

with *ϕ* := 0.5, which is the state-expectation parameter,
- *p*(*c*_*t*_|*s*_*t*_) is the state-conditional contrast-difference distribution defined by the uniform distribution,

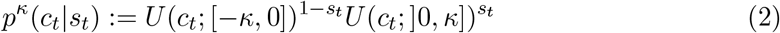
- 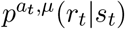 is the action- and contingency-parameter-dependent and state-conditional reward distribution. This distribution is defined by

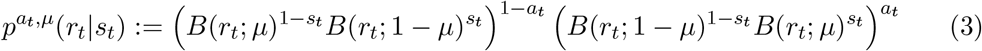

with contingency parameter *µ* := 0.9 for half of the blocks and *µ* := 0.1 for the other half under lower reward uncertainty. Similarly, the contingency parameter *µ* := 0.7 for half of the blocks and *µ* := 0.3 for the other half under higher reward uncertainty.

### Gabor-Bandit agent model

In our computational model, we assumed three computational stages corresponding to perceptual inference modeling visual processing of the displayed information, learning about the reward probabilities, and economic decision-making.

### Perceptual inference

To model perceptual inference, we assume

- *O∈* ℝ is the set of the agent’s internal observations *o*_*t*_ that are dependent on the contrast difference of the external Gabor patches *c*_*t*_ along with perceptual uncertainty or noise,
- 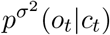 is the agent’s observation likelihood, defined as the contrast difference-conditional observation distribution,

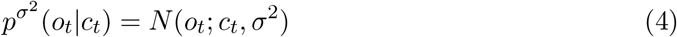

where, in our simulations, we manipulate *σ* to induce high (*σ* = 0.03) and low (*σ* = 0.0001) levels of perceptual uncertainty.

To compute the agent’s belief state dependent on the observed contrast difference, we have

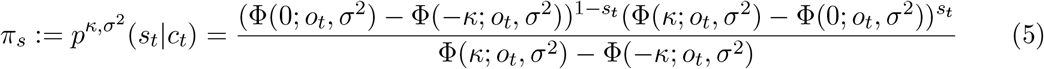

where Φ is the Gaussian cumulative distribution function (CDF).

### Economic decision-making

For economic choices, we considered the following variables.

- In the agent, *µ* is a random variable representing the contingency parameter,
- M := [0,1] is the outcome space of this random variable,
- *p*(*µ*) is the agent’s belief about the task-block contingency parameter

We assumed that the agent model chooses action 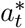 with the higher expected reward

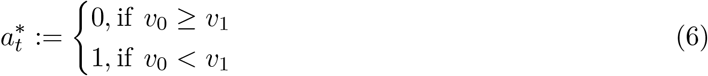

where the expected value conditional on action *a*_*t*_ = 0 is given by

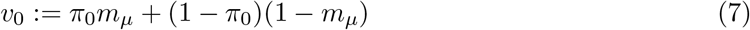

and conditional on action *a*_*t*_ = 1 by

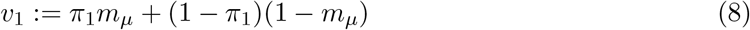

Where

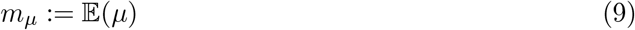

is the average of the contingency parameter.

### Learning

To learn from the presented reward feedback, the agent updates the distribution over the contingency parameter

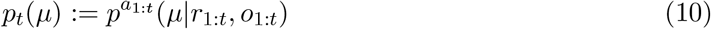

This is achieved by evaluating the polynomials in *µ*, where the polynomial coefficients *ρ*_*t*,0_, …, *ρ*_*t,t*_ of

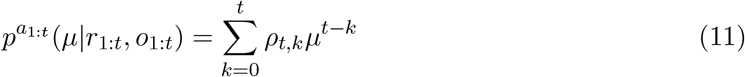

can be evaluated based on *ρ*_*t−*1,0_, …, *ρ*_*t−*1,*t−*1_ of

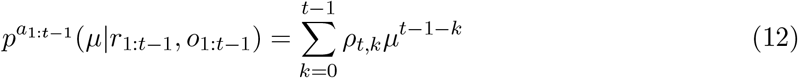

where

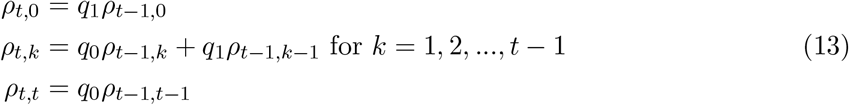

and where

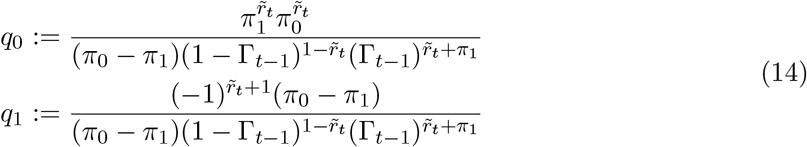

With

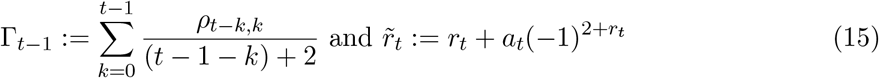

### Data preprocessing

For our statistical analyses, we relied on participants’ single-trial slider responses, from which we derived updates, prediction errors, and learning rates.

- 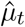 indicates the subject’s slider response, which we take to indicate the subject’s belief about the contingency parameter *µ*_*t*_ in the Gabor-Bandit task. Please recall that we used congruent (the subject was asked to report the contingency parameter of the “correct”, i.e., more rewarding option) and incongruent (the subject was asked to report the contingency parameter of the “incorrect”, i.e., less rewarding option) blocks in our experiment. To map the slider responses on congruent and incongruent blocks onto a common scale, we recoded responses on incongruent blocks according to

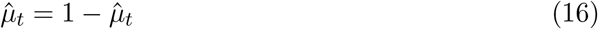
- *Q ∈ {*0, 1*}* indicates a correct (*q* = 1) and incorrect choice (*q* = 0), defined by

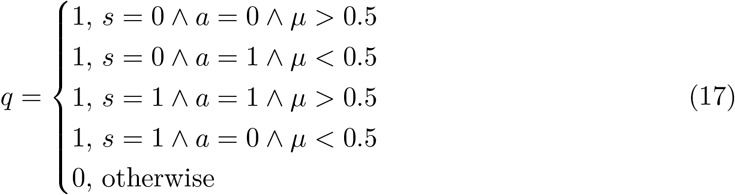
- *D ∈* [*−*1, 1] denotes the set of prediction errors, defined by

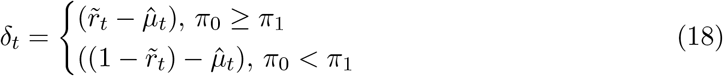

where 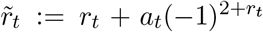. That is, when computing the prediction error, we take into account the state-action-reward contingency defined in the task model (eq. (3)). For example, when the presented contrast difference favors state *s*_*t*_ = 0, we assume *π*_0_ *> π*_1_ and conditional on action *a*_*t*_ = 0, the expected reward probability is 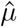. To account for the action dependency of the reward, we rely on 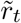, so that, for example, *r*_*t*_ = 0 conditional on *a*_*t*_ = 1 corresponds to 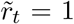 re-coded with respect to action *a*_*t*_ = 0 (where *r*_*t*_ = 1 had it been chosen). Similarly, to account for the state dependency of the reward, we rely on (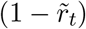) when state *s*_*t*_ = 1 is more likely than *s*_*t*_ = 0,
- *U ∈* [*−*1, 1] denotes the set of updates, defined by

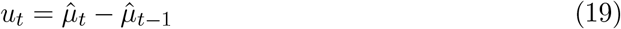
- *B ∈*{0, 1} indicates a choice-confirming outcome (*b*_*t*_ = 1) and a choice-dis-confirming outcome (*b*_*t*_ = 0), defined by

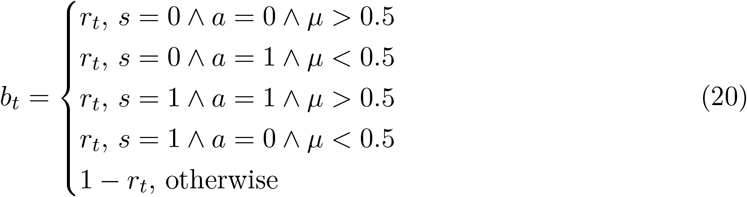
- *K ∈*{0, 1} denotes the set of congruence-trial types, where *k*_*t*_ = 0 denotes an incongruent and *k*_*t*_ = 1 a congruent trial type,
- *L ∈*{0, 1} denotes the set of salience-trial types, where *l*_*t*_ = 0 denotes a low-salience and *l*_*t*_ = 1 a high-salience trial.

### Regression analysis

To better understand the factors influencing the single-trial updates, we used a regression model that allowed us to dissociate multiple factors driving the learning rate. This regression model can be interpreted through the lens of reinforcement learning, according to which prediction errors, scaled by a learning rate, determine belief updates (Daw, 2014; McGuire et al., 2014):

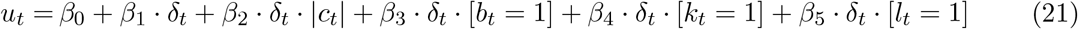

*β*_0_ is the intercept. *β*_1_ is the coefficient modeling the average effect of the prediction error on the update, which we interpret as the fixed learning rate, as common in reinforcement learning (usually denoted *α*). We refer to this term as fixed LR. To model how flexibly participants adjusted their learning for belief states emerging from various levels of contrast differences, we added the interaction term *β*_2_ between prediction error and absolute contrast difference. We refer to this term as belief-state-adapted LR. Please note that we excluded trials from the reward-uncertainty condition as contrast differences on these trials were high, and hence perceptual uncertainty was not induced. Next, to check for the presence of confirmation biases in learning, we use the interaction term *β*_3_ between prediction error and whether an outcome is confirming (confirmation bias). This is coded as a categorical variable, i.e., 0 for outcomes that dis-confirm the choice and 1 for outcomes that confirm the choice. Finally, we added two task-based block-level categorical variables as control regressors. *β*_4_ was the interaction term between salience (high vs. low contrast blocks) and prediction error where 0 denoted trials in a low-contrast block and 1 for trials in a high-contrast block, and *β*_5_ captured effects of congruence (congruent vs. incongruent block type) in interaction with prediction error where 0 denoted trials in an incongruent block and 1 for trials in a congruent block. All continuous regressors, except for prediction errors, were re-scaled within the range of 0 and 1 using the min-max normalization method

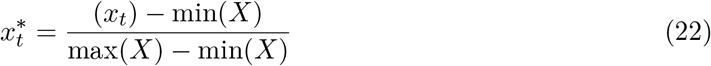

where *X* is the variable of interest, *x*_*t*_ is the value on a given trial that gets normalized, and 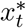 is the normalized value for a given trial. For prediction errors, we used its natural scale since it was key to retain its valence for the signed LR analyses.

The model was fit to each participant’s single-trial updates. Since prediction errors *δ*_*t*_ = 0 do not call for learning, we excluded such trials. Moreover, one potential drawback of using a canonical linear regression model is the assumption that the residuals are homoscedastic, that is, similar across the range of the predictor variable. However, in our model, the assumption of homoscedasticity is violated, particularly for larger prediction errors. Thus, we accounted for heteroscedasticity by using a weighted regression model, wherein more weight is given to the observations with smaller residuals providing more reliable information.

### Regression diagnostics

We used two statistical tools to illustrate the regression coefficients. First, to illustrate the incremental effect of a specified regressor on the single-trial updates, after accounting for the effects of all other terms, we created a partial regression plot (also known as an added variable plot). This plot is formed by plotting the (i) residuals from regressing single-trial updates against all regressors except the regressor of interest versus (ii) residuals from regressing the specified regressor against all the remaining regressors. This type of analysis emphasizes the marginal contribution of a given regressor in capturing the participant’s updates over and above all the other regressors. Second, we used an interaction plot to demonstrate the dynamics of interaction regressors on single-trial updates. We plotted the conditional effect of prediction errors given specific values of the other task-based variable in the interaction. For categorical regressors, the specific values were set to the different categories of the variable. For continuous regressors, we used three values, each corresponding to the lowest, highest, and median values. To plot this, we compute the adjusted model-predicted update for an observation of all the regressors contributing to an interaction term while averaging out the effect of the other regressors (also known as adjusted response).

## Data and code availability

All data and code will be made available on GitHub at the time of publication.

## Acknowledgements

We thank Hauke Heekeren for his mentorship and amazing support throughout the project. We also thank Muhammad Hashim Satti for helpful comments on an earlier draft of the manuscript. P.G. was supported by Deutscher Akademischer Austauschdienst (DAAD) Graduate School Scholarship Programme, 2020. R.M.C. was supported by The German Research Council grants (CI241/3-1, INST 272/297-1) and the European Research Council grant (ERC-StG-2018-803370). N.W.S. is funded by a Starting Grant from the European Union (ERC-2019-StG REPLAY-852669) and the Federal Ministry of Education and Research (BMBF) and the Free and Hanseatic City of Hamburg under the Excellence Strategy of the Federal Government and the Länder. C.F. was supported by German Research Foundation (DFG), grant number FI 2309/1-1. R.B. was supported by DFG (Deutsche Forschungsgemeinschaft) grant 412917403.

## Supplementary material

### Extended results

#### Absolute learning-rate analysis

Our analysis of signed learning rates based on the regression models shows that prediction errors in conjunction with multiple factors, such as belief states and choice-confirming outcomes, govern learning rates. However, one potential issue of our signed learning-rate approach is that lower learning rates could be an indicator of (i) a strategic calibration of the learning rate to perceptual uncertainty or (ii) more frequent confusion of the task states due to perceptual uncertainty. The first interpretation (strategic adjustment of the learning rate) would be in line with our hypothesis that humans adjust learning to uncertainty. However, according to the second interpretation (state confusion), lower fixed and belief-state-driven learning rates would arise when subjects misperceive the stimuli and learn in the wrong direction. To tease these two interpretations apart, we analyzed absolute prediction errors and updates. Running the analyses on absolute prediction errors and updates yields learning-rate estimates of how much participants learned independently of whether they learned in the correct or incorrect direction. As such, this approach allows us to examine the magnitude of updates independent of whether they confused the task states or not.

In line with the perspective that learning behavior is shaped by prediction errors, we found a significant correlation between absolute prediction errors and updates (Fig. S1a). Additionally, we found a significant relationship between contrast differences and absolute single-trial updates (Fig. S1b). That is, participants made larger updates on the slider when the contrast difference was larger.

However, the single-trial approach might also be more strongly affected by response noise, and we, therefore, next applied our regression model to absolute prediction errors and updates. The fixed learning rate reflecting the average influence of prediction errors on absolute updates was positive (mean = 0.13 *±* 0.02, *t*_97_ = 6.31, *p <* 0.001, Cohen’s *d* = 0.64) (Fig. S1c, fixed-LR coefficient). This confirms our results from the analysis of signed learning rates that prediction errors drive learning.

Additionally, contrast differences appear to have a similar influence on absolute and signed learning. Consistent with the signed learning-rate approach, we found that larger contrast differences propelled absolute updates for a given prediction error, as indicated by the positive coefficients for belief-state-adapted learning (mean = 0.05 *±* 0.014, *t*_97_ = 3.47, *p <* 0.001, Cohen’s *d* = 0.35) (Fig. S1c, belief-state-adapted-LR coefficient). This implies that absolute updates increase with increasing contrast-difference levels for a given prediction error (Fig. S1d; example participant). Additionally, across participants, signed belief-state-adapted-LR coefficients were strongly correlated with absolute coefficients (Fig. S1e), suggesting that both approaches capture dynamic learning in a comparable way.

Finally, we found evidence of the confirmation bias, similar to the signed learning-rate analysis. In our regression model, positive confirmation-bias coefficients indicate stronger updates following outcomes that confirm the participant’s choice (mean = 0.1 *±* 0.009, *t*_97_ = 10.61, *p <* 0.001; Cohen’s *d* = 1.07 (Fig. S1c, confirmation bias; Fig. S1f). Again, these results coincide with the impact confirming outcomes had in the signed learning-rate approach.

**Figure S1.**
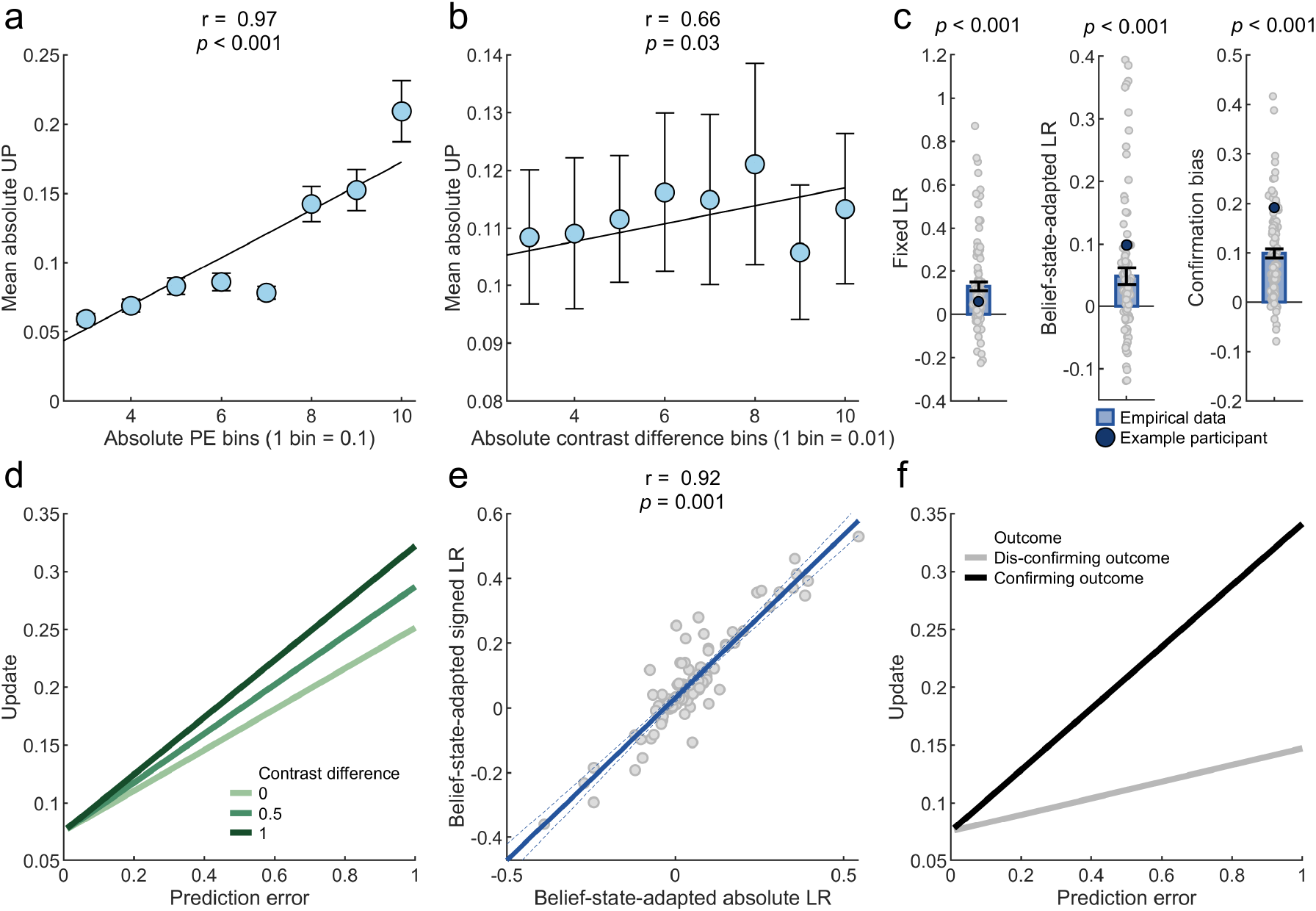
Absolute learning-rate analysis. **a**| Mean *±* standard error of the mean (SEM) absolute single-trial updates grouped across 10 absolute single-trial prediction-error bins (Pearson’s *r*_08_ = 0.97, *p* < 0.001). Participants’ slider updates were larger for larger prediction errors. **b**| Mean *±* SEM absolute single-trial updates grouped across 10 contrast-difference bins (Pearson’s *r*_08_ = 0.66, *p* = 0.038). **c**| Mean *±* SEM coefficients for key regressors from the linear regression model fit to absolute single-trial updates. Positive fixed-LR coefficients indicate participants’ proclivity to show larger updates for larger absolute prediction errors (Cohen’s *d* = 0.64). Similarly, belief-state-adapted-LR coefficients convey a contrast-difference-contingent update magnitude (Cohen’s *d* = 0.35). The confirmation-bias coefficient also revealed higher absolute learning from confirming outcomes (Cohen’s *d* = 1.07). **d**| Across three levels of contrast-difference values, regression fits for a range of absolute prediction-error values show contrast-difference-modulated flexible learning. Higher contrast differences led to larger updates, presumably driven by more distinct belief states, as compared to lower contrast differences, for a given prediction error. **e**| Relationship between absolute and signed belief-state-adapted LR across participants shows that both approaches to analyzing the data corroborate the presence of flexible learning (Pearson’s *r*_08_ = 0.92, *p* = 0.001). **f**| Larger updates were made on trials where the participant learned from outcomes that confirmed the participant’s belief estimate, across different values of prediction errors.

#### Extended learning-rate analysis

Next to adjustments in learning rates for prediction errors and belief states (see eq. (21)), we now discuss the impact of choice confirmation and additional control regressors on signed updates. We found that choice-confirming outcomes impacted signed and absolute updates. These findings align with existing literature showing higher learning rates for choice-confirming outcomes, as compared to negative or neutral information that dis-confirms choices (Lefebvre et al., 2017; Nickerson, 1998; Palminteri et al., 2017; Pupillo & Bruckner, 2023; Sharot & Garrett, 2016). Studies suggest that the bias can be potentially beneficial in a risky environment in which outcomes can only partially be predicted. Learning preferentially from choice-confirming outcomes might yield more robust expected-value representations since dis-confirming outcomes that are due to outcome variability hold less sway over expected values (Kandroodi et al., 2021; Lefebvre et al., 2017; Palminteri & Lebreton, 2022; Tarantola et al., 2021). Crucially, based on our study, it is not possible to clearly dissociate the confirmation bias (stronger learning from choice-confirming outcomes) from the positivity bias (stronger learning from positive outcomes), which might require a comparison between instrumental (as in our task) and Pavlovian tasks (where due to the absence of choices, only the positivity bias can show up; Lefebvre et al. (2017)).

Furthermore, the contrast (higher vs. lower) of the more rewarding option in a block termed as the “salience” of the more rewarding patch (mean = *−* 0.01 *±* 0.009, *t*_97_ = *−* 1.52, *p* = 0.13, Cohen’s *d* = 0.15) and slider congruence (congruent vs. incongruent) (mean = 0 ± 0.009, *t*_97_ = *−* 0.15, *p* = 0.88, Cohen’s *d* = 0.01) did not have a significant effect on updates. Similarly, these regressors did not have a significant impact (mean = *−* 0.02 *±* 0.009, *t*_97_ = *−* 1.8, *p* = 0.08, Cohen’s *d* = 0.18; Salience and mean = 0 *±* 0.009, *t*_97_ = *−* 0.19, *p* = 0.85, Cohen’s *d* = 0.02; Congruence) on absolute updates (Fig. S2a). In addition to this result being in line with the normative agent, this also clarifies that peripheral task factors did not impact learning.

**Figure S2.**
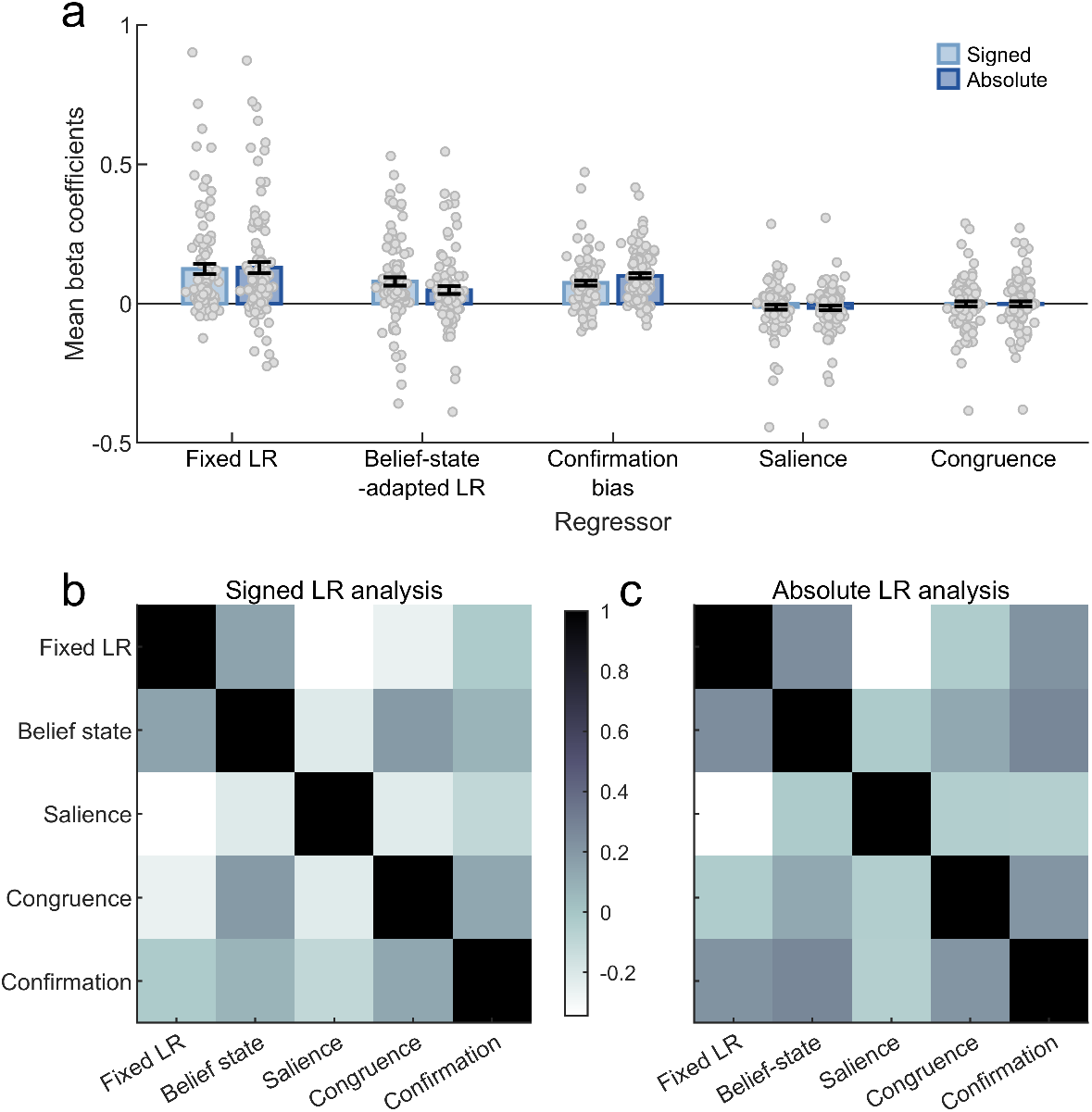
Full regression model and multi-collinearity check. **a**| Mean *±* standard error of the mean (SEM) coefficients for all key and control regressors from the signed and absolute linear regression model. **b**| Heat-map showing correlation coefficients between coefficient values for all regressors from the signed learning-rate analysis. **c**| Heat-map showing correlation coefficients between coefficient values for all regressors from the absolute learning-rate analysis.

Additionally, we also checked for correlations between the estimated coefficients for both signed and absolute analysis. Correlation matrices show correlations in the low-to-moderate range between the estimated coefficients for both sets of analysis (Fig. S2b-c). This indicates that estimated coefficients are not spuriously exaggerated or mitigated, which could, in principle, be a result of multi-collinearity.

Moreover, to control for how learning changed with the different levels of reward probability, we added an additional term modeling the interaction between prediction error and the level of reward uncertainty (risk-adapted LR) as a control regressor to eq. (21). This is coded as a categorical variable, i.e., 0 for high reward uncertainty and 1 for low reward uncertainty. This control analysis yielded that risk did not significantly impact signed (mean = *−* 0.02 *±* 0.019, *t*_98_ = *−* 1.31, *p* = 0.2, Cohen’s *d* = 0.15) and absolute (mean = *−* 0.01 *±* 0.018, *t*_97_ = *−* 0.62, *p* = 0.72, Cohen’s *d* = 0.06) updates (Fig. S3a). However, risk-adapted LR coefficients were correlated with fixed LR (Pearson’s *r*_97_ = *−* 0.62, *p <* 0.001), and we, therefore, decided to exclude it from the regression model.

**Figure S3.**
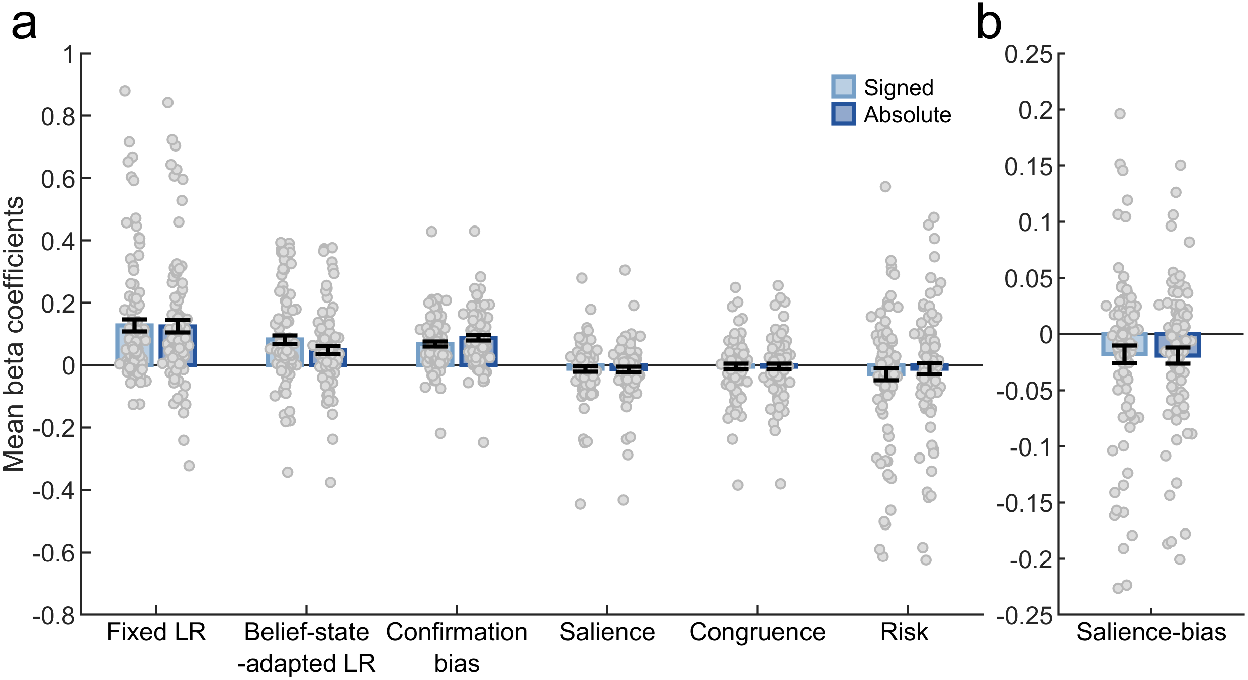
Full signed and absolute learning-rate analyses. **a**| Mean *±* standard error of the mean (SEM) coefficients for all key and control regressors, including risk-adapted learning-rate coefficients from the signed and absolute linear regression model. **b**| Mean *±* SEM coefficients for the salience-bias coefficient.

We also extended the regression model to clarify if the salience bias identified during decision-making impacts learning. We added this as an interaction term between prediction errors and a categorical variable representing salience (low vs. high), which denotes if the more or less salient option was chosen on the given trial. Negative significant coefficients for this regressor show that participants preferentially up-regulated signed (mean = *−* 0.02 *±* 0.008, *t*_97_ = *−*2.28, *p <* 0.05, Cohen’s *d* = 0.23) and absolute (mean = *−* 0.02 *±* 0.007, *t*_97_ = *−* 2.7, *p <* 0.001, Cohen’s *d* = 0.27) updates after choosing the less salient option (Fig. S3b). This effect could reflect a strategy to compensate for the lower subjective expected value due to the salience bias affecting economic decision-making (via stronger prediction errors).

#### Model validation

**Figure S4.**
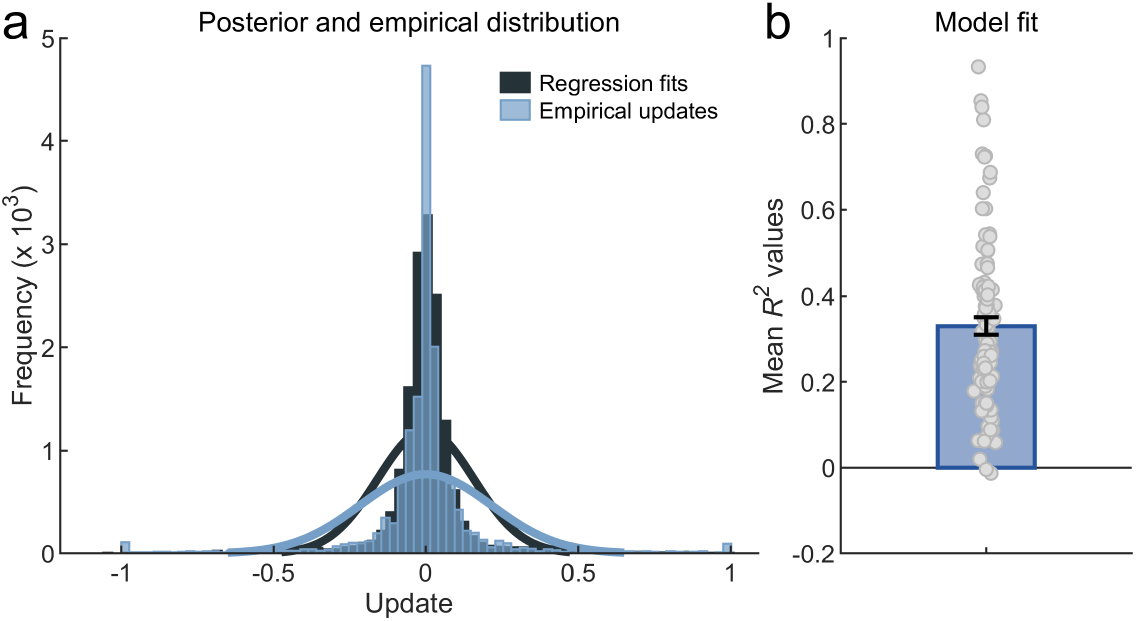
Model-fit assessment. **a**| A visual representation of the goodness of fit, as illustrated by the model-predicted posterior updates using estimated parameters and single-trial regression data. **b**| *R*^2^ values show the regression model was moderately effective in capturing and explaining learning data despite heterogeneity across participants.

To systematically compare the regression results to the empirical data, we performed posterior-predictive checks. Model-based updates captured the general trend in participants’ updates (Fig. S4a). One key difference is that empirical updates included a high frequency of extremely small updates (around 0 as indicated by the blue bar in Fig. S4a). We identified trial-by-trial variation in motor noise while responding with the slider as one potential reason for these extremely small updates. Such empirical updates are regardless of prediction errors and task-based variables. The regression fits feature extremely small posterior updates to a lesser extent since posterior updates are systematically scaled depending on the prediction error and other predictors of the model on a given trial.

We also illustrate that the model captures the learning dynamics for individual participants who also show evidence for a higher frequency of updates around 0 (Fig. S5). Finally, we examined the regression model’s goodness of fit using *R*^2^, suggesting a moderate fit (Fig. S4b).

**Figure S5.**
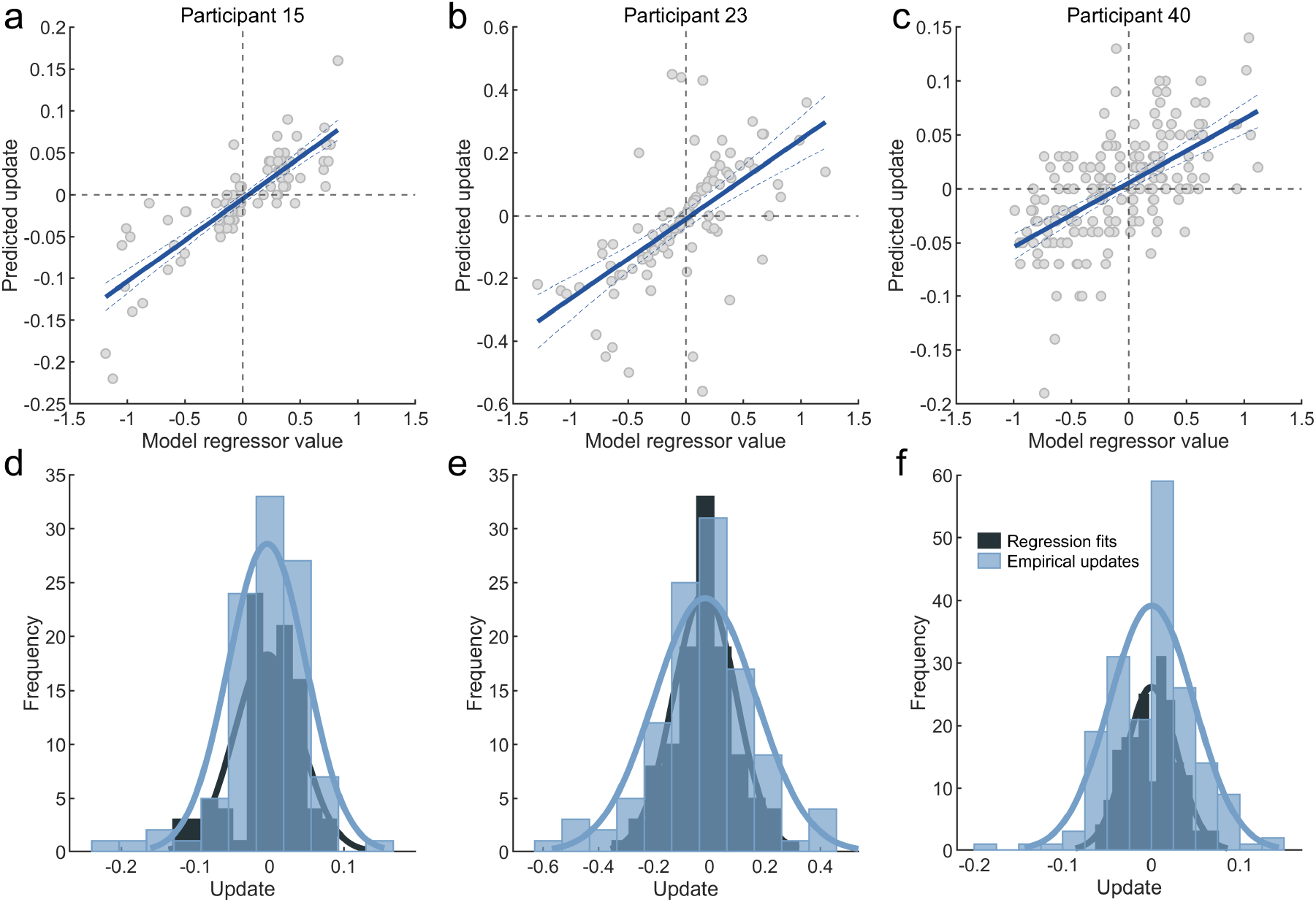
Example participant diagnostics. **a-c**| Added variable plots assessing the relationship between all model regressors and updates. The evident linear relationship (dark blue line) suggests that the model regressors made impactful contributions in capturing the general trend of single-trial updates for individual participants. **d-f**| Single-subject posterior updates predicted by the model efficiently capture single-subject updates.

**Figure S6.**
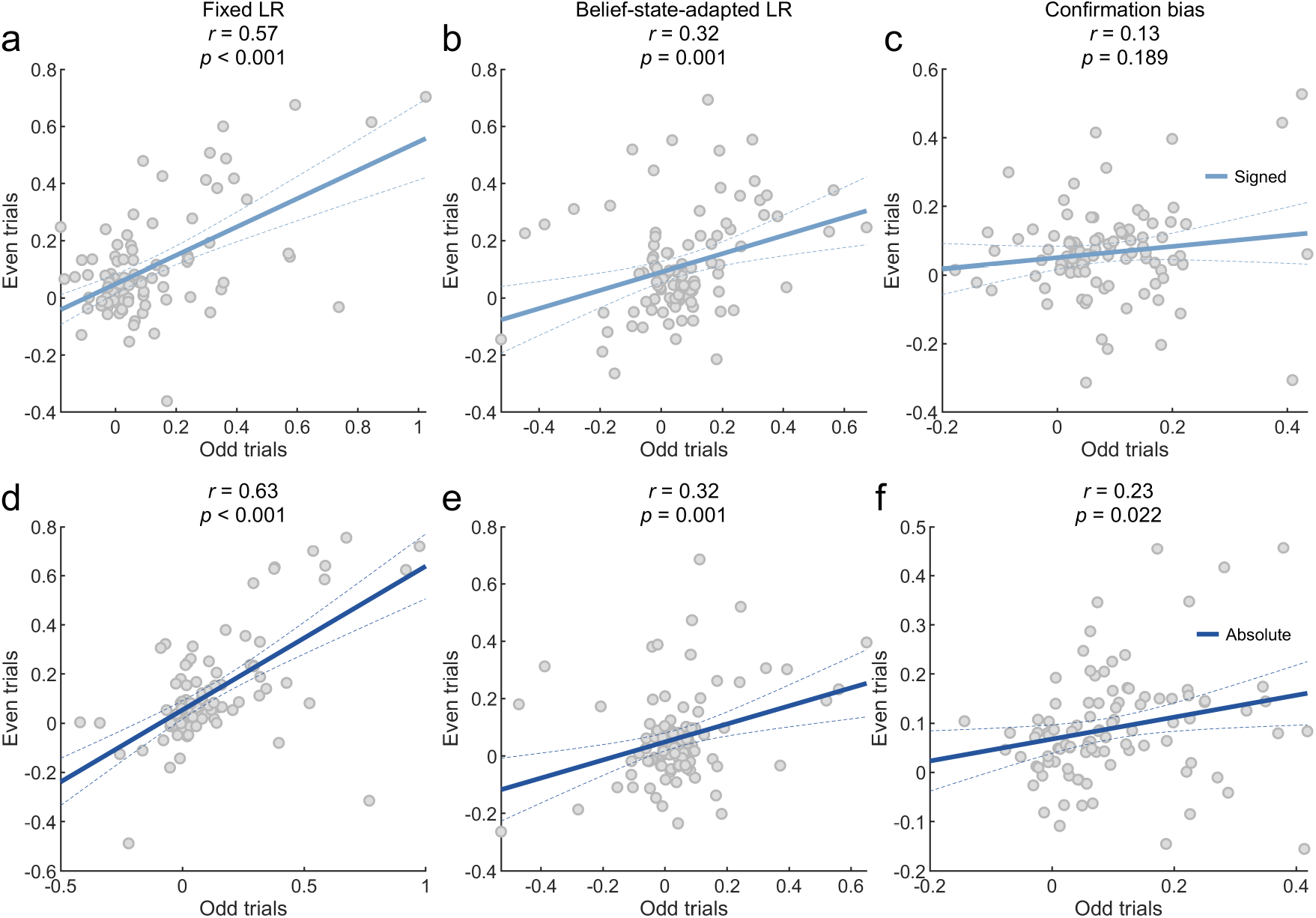
Split-half reliability correlation coefficients between odd and even trials. Correlation between signed-analysis coefficients for **a**| fixed LR, **b**| belief-state-adapted LR, and **c**| confirmation bias. Correlation between absolute-analysis coefficients for **d**| fixed LR, **e**| belief-state-adapted LR, and **f**| confirmation bias.

### Split-half reliability for fixed and flexible learning parameters

To test how internally consistent our model’s estimated fixed and flexible learning parameters were, we adopted the split-half reliability measure. This involved grouping odd and even trials into different sub-sets to run separate regressions to obtain fixed and flexible learning rate coefficients for each subset. To quantify reliability, we computed the Pearson’s correlation coefficient between the parameters estimated (Fig. S6). We found that fixed learning-rate coefficients have moderate reliability (Pearson’s *r*_98_ = 0.57, *p <* 0.001; signed analyses and Pearson’s *r* = 0.63, *p <* 0.001; absolute analyses). However, we found weaker reliability measures for both belief-state-adapted LR (Pearson’s *r*_98_ = 0.32, *p <* 0.01; signed analyses and Pearson’s *r*_98_ = 0.32, *p <* 0.01; absolute analyses) and confirmation bias (Pearson’s *r*_98_ = 0.13, *p* = 0.19; signed analyses and Pearson’s *r*_98_ = 0.23, *p <* 0.05; absolute analyses).

### Extended belief-accuracy analysis

**Figure S7.**
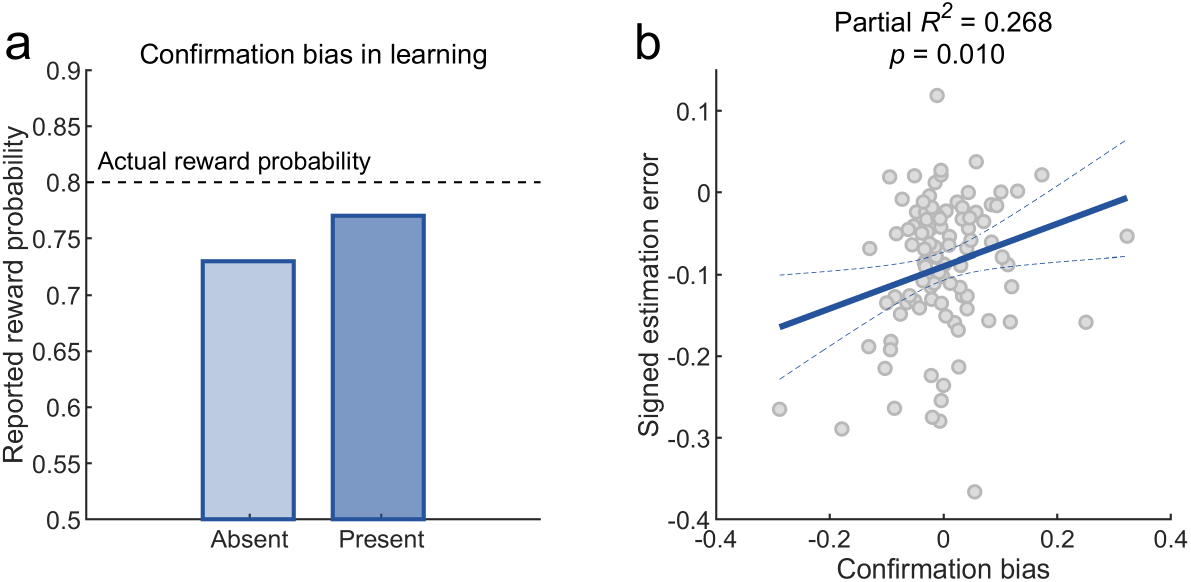
Confirmation bias and signed estimation error. **a**| Illustration of the hypothetical role of the confirmation bias during learning under uncertainty. The confirmation bias reflects stronger learning from choice-confirming than dis-confirming outcomes. In some situations, it might boost learned value representations. In this example, the confirmation bias helps the learner estimate the value more accurately (lower underestimation of the true value) compared to an unbiased learner (higher underestimation of the true value). **b**| We tested this idea based on the experimental data. To do so, we relied on signed estimation errors indicating the degree of under-versus overestimation of the true but unknown reward probability. Most subjects tended to underestimate the reward probability (average estimation error across all blocks). The confirmation bias and the signed estimation error turned out to be associated in that stronger confirmation biases statistically predicted more accurate reward probabilities (less underestimation of the true probabilities). This result is consistent with the idea that under some circumstances, the confirmation bias can be adaptive.

#### Confirmation bias and over-estimated beliefs

To empirically test whether the confirmation bias is linked to the extent to which participants under-or overestimated the actual contingency parameter, we examined the relationship between our regression coefficients and signed estimation errors. We quantified signed estimation errors as the signed difference between the actual reward contingency and the value reported by the participant, which corresponds to whether participants over-or underestimated the reward probability. Due to reward uncertainty in our task, correct choices were sometimes not rewarded (e.g., in 30%, correct choices were not rewarded). It has been argued that a confirmation bias is beneficial to learning under such challenging conditions: When choice-confirming outcomes have a stronger effect on the learning rate, value representations might become more robust and might potentially be more weakly affected by reward uncertainty (Lefebvre et al., 2017; Palminteri et al., 2017). This could result in overestimated reward probabilities compared to an unbiased strategy (Fig. S7a). Indeed, we found a significant relationship between signed estimation error and confirmation bias (*β* = 0.25, *p <* 0.01; Fig. S7b). That is, participants with higher confirmation bias showed less negative estimation errors, suggesting that the confirmation bias might have helped them calibrate learning to reward uncertainty. In line with this perspective, we found that participants with stronger choice-confirmation biases tended to show reduced underestimation of the actual reward probability.

**Figure S8.**
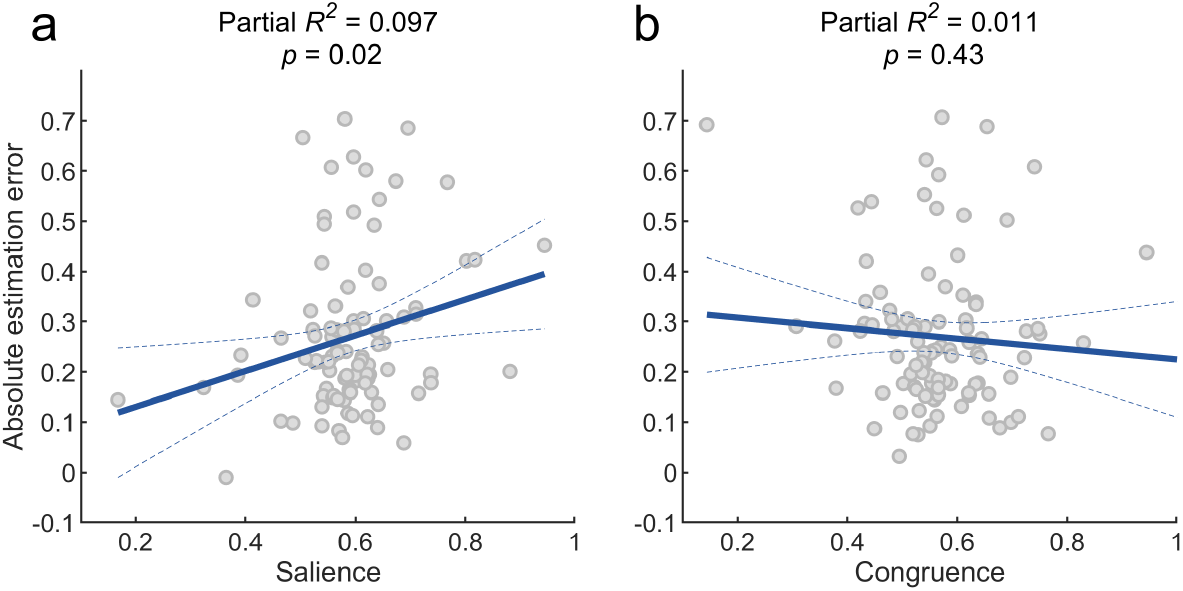
Influence of learning on belief accuracy. Relationship between absolute estimation error and **a**| salience and **b**| congruence.

#### Signed learning rate and belief accuracy

Next, we also controlled for potential links between the adapted signed learning rate for control regressors from equation (21) and absolute estimation error. Salience had a significant impact on estimation error (*β* = 0.36, *p* = 0.02) whereas congruence did not have a significant relationship with estimation error (*β* = *−* 0.1, *p* = 0.428) (Fig. S8).

**Figure S9.**
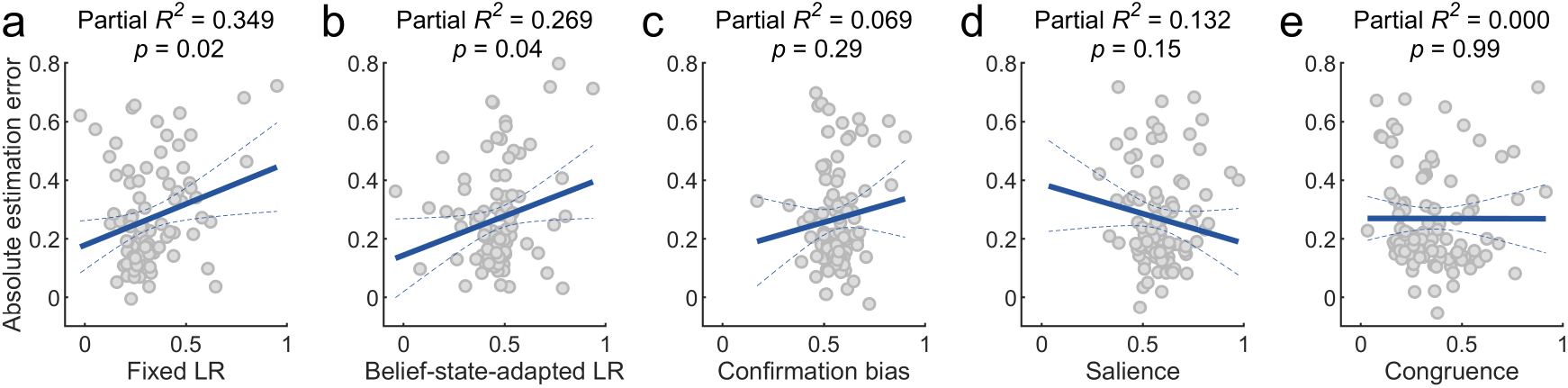
Influence of absolute learning rates on belief accuracy. Relationship between absolute estimation error and coefficients for **a**| fixed LR, **b**| belief-state-adapted LR, **c**| confirmation bias, **d**| salience, and **e**| congruence.

#### Absolute learning rate and belief accuracy

To check if absolute learning rates impacted belief accuracy, we fit a model containing all learning-rate coefficients from the absolute learning-rate analyses to absolute estimation errors. We found that subjects with high absolute fixed learning-rate coefficients (i.e., prediction-error-driven learning) tended to have larger estimation errors (*β* = 0.28, *p* = 0.02; Fig. S9a). Similarly, individual differences in belief-state-adapted-LR coefficients had a significant relationship with estimation error (*β* = 0.267, *p* = 0.04; Fig. S9b). We found no significant links between estimation error and confirmation bias LR (*β* = *−* 0.0014, *p* = 0.989; Fig. S9c). Similarly, we found no significant links between estimation error and salience (*β* = 0.199, *p* = 0.293; Fig. S9d). Finally, we found that congruence did not significantly impact estimation errors (*β* = *−*0.2028, *p* = 0.147; Fig. S9e).

#### Pilot study

We used a reduced version of the current task design (excluding the slider) in a pilot study. We integrated a combination of both perceptual and reward uncertainty as an extension to the Gabor-Bandit task (Bruckner et al., 2020). We collected pilot data of 100 participants (52 female, 48 male; mean age = 22.91 *±* 3.04; age range 18-30). We excluded data from seven participants as they performed with less than 50% accuracy. Participants completed a total of 16 blocks, with 25 trials each. Each block belonged to one of three within-subject experimental conditions similar to the Experimental task. We also added a fourth control condition with low levels of perceptual and reward uncertainty. For conditions with high perceptual uncertainty, we sampled contrast differences from [−0.08, 0] when the left patch had the lower contrast (*s*_*t*_ = 0) and [0, 0.08] when the right patch had the lower contrast (*s*_*t*_ = 1). In the conditions with low perceptual uncertainty, the contrast difference was in the range [−0.38, −0.3] when the contrast of the left patch was lower (*s*_*t*_ = 0) and [0.3, 0.38] when the contrast of the right patch was lower (*s*_*t*_ = 1). Finally, we counterbalanced the mapping between states, actions, and rewards. The order of the conditions was randomized for each participant. However, for the first fifty participants, the order was not completely randomized. The first and the eighth blocks deterministically belonged to the both-uncertainties condition.

#### Replicating decision-making results in pilot study

Results from the pilot study align with the salience bias seen in the analysis of choices from the primary Experimental task. Participants showed a significant salience bias in the both-uncertainties condition (mean = 0.07 *±* 0.017, *t*_92_ = 4.26, *p <* 0.001), reward condition (mean = 0.09 *±* 0.016, *t*_92_ = 5.71, *p <* 0.001), and control condition (mean = 0.04*±*0.008, *t*_92_ = 5.65, *p <* 0.001). However, in the perceptual-uncertainty condition (mean = 0.02*±* 0.013, *t*_92_ = 1.22, *p* = 0.22), we did not find a significant salience bias. Next, we tested if the salience bias is more enhanced due to reward uncertainty. Participants showed a significantly pronounced salience bias in the both-uncertainties condition as compared to the perceptual-uncertainty condition (*t*_92_ = 3.25, *p <* 0.01, Cohen’s *d* = *−* 0.38). Participants showed a significantly larger salience bias in the no-uncertainty condition as compared to the reward-uncertainty condition (*t*_92_ = 3.07, *p <* 0.001, Cohen’s *d* = 0.41) (Fig. S10a). We also fitted a linear regression model to the participants’ average economic performance in a block. We used regressors that corresponded to the (i) perceptual (salience and uncertainty) and (ii) reward information on the block level (Fig. S10b). Crucially, in support of the hypothesis that humans integrate value and salience, positive coefficients for the main effect of salience reveal that economic choices were significantly more likely to be correct on high-contrast blocks, as opposed to low-contrast blocks (main task: mean = 0.06 *±* 0.014, *t*_97_ = 4.4, *p <* 0.001, Cohen’s *d* = 0.44, pilot study: mean = 0.06*±* 0.009, *t*_92_ = 6.49, *p <* 0.001; Cohen’s *d* = 0.67). Additionally, a negative and significant coefficient for high perceptual uncertainty shows that participants performed worse on blocks with high perceptual uncertainty, as compared to low perceptual uncertainty (main task: mean = *−* 0.07 *±* 0.013, *t*_97_ = −5.61, *p <* 0.001, Cohen’s *d* = 0.57, pilot study: mean = *−* 0.09*±* 0.009, *t*_92_ = −9.51, *p <* 0.001; Cohen’s *d* = 0.99). Finally, we also found that economic choice performance was significantly worse in blocks where reward uncertainty was high, as captured by negative coefficients for the main effect of high reward uncertainty (main task: mean = *−*0.13 *±* 0.012, *t*_97_ = −11.29, *p <* 0.001, Cohen’s *d* = 1.14, pilot study: mean = *−*0.18 *±* 0.012, *t*_92_ = −15.16, *p <* 0.001; Cohen’s *d* = 1.57).

**Figure S10.**
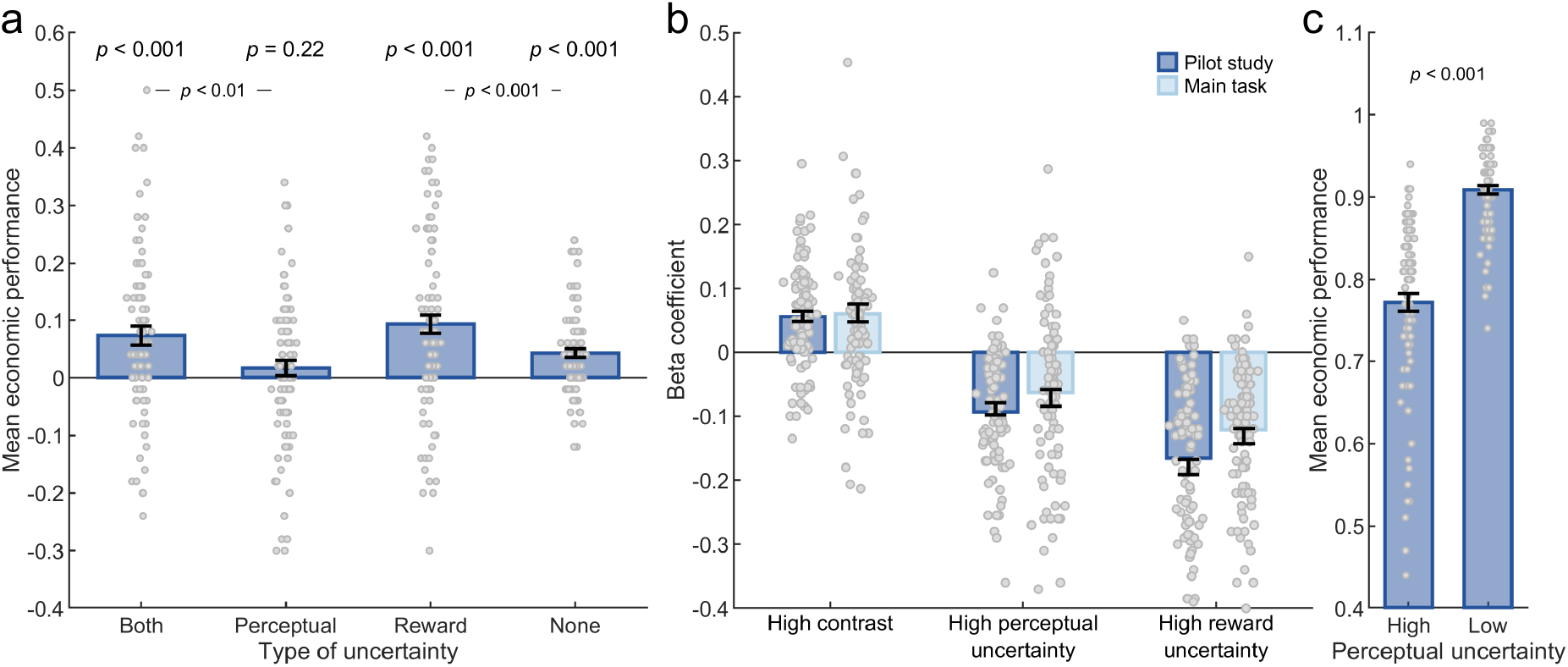
Choice analysis of pilot experiment. **a**| Mean*±* standard error of the mean (SEM) salience bias for types of uncertainties. Positive salience bias indicates participants’ preference for the high-salience option. **b**| Mean ± SEM coefficients for key regressors after fitting a linear regression model to block-level choice accuracy. Positive coefficients for the main effect of high contrast indicate participants’ proclivity to choose the high salience option. Negative coefficients for high levels of perceptual and reward uncertainty capture the decrease in participant’s performance with increasing uncertainty in the environment. **c** Mean *±* standard error of the mean (SEM) choice performance across levels of perceptual uncertainty showing that high perceptual uncertainty leads to worse performance.

**Figure S11.**
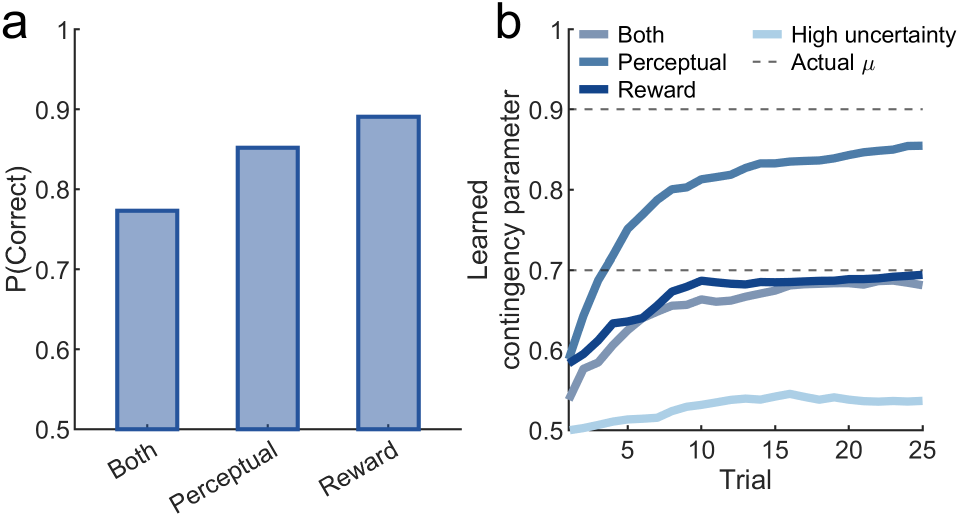
Normative agent’s simulated choice and learning behavior. **a**| Averaged across 100 simulations, choice performance is corrupted by higher perceptual uncertainty (“both” and “perceptual” condition). **b**| Learned contingency parameter (*µ*) converges towards the actual contingency parameter. Simulated learning curves are more noisy in the higher reward uncertainty blocks due to riskier outcomes that the agent is required to learn from. In comparison to blocks with low reward uncertainty, the agent shows systematically slower learning in the higher reward uncertainty blocks (“both”, “reward” and “high uncertainty” condition). Additionally, we see slower learning due to extremely high perceptual uncertainty which primarily dictates the agent’s learning patterns (see learning curve for high-uncertainty blocks) in conjunction with reward uncertainty.

## Extended task details

### Practice task

Before taking part in the main task, participants were trained using an adapted version. The specific details differed across both studies.

**Study 1** Participants performed four practice blocks, with 50 trials each. On half of the practice blocks, participants were presented with high perceptual uncertainty trials. However, reward-uncertainty levels were not manipulated across the two practice blocks. Thus, the latent state-action-reward contingency was such that on half of the practice blocks, the patch with higher contrast had a 100% reward probability, while on the other half, the patch with lower contrast had a 100% reward probability. The trial structure was the same as the main task and participants were expected to make an economic decision. The main aim of the practice blocks was to train participants in an easier version of the main task.

**Study 2** Participants performed three practice blocks, with 25 trials each. Each block belonged to each of the three uncertainty conditions. Specifically, the blocks were sequentially presented to ensure increasing order of difficulty. Participants started off with the perceptual uncertainty condition followed by the reward uncertainty condition. Finally, participants did the both-uncertainties condition. The trial structure was the same as the main task and participants were expected to make an economic decision and learn to use the slider to report their estimated contingency parameter.

### Instructions

Participants were presented with an online version of the instructions. Multiple images demonstrated various stages of the task which was accompanied by written explanation. Post this, participants were asked to answer questions about the task in a quiz. For every incorrect answer, participants were reminded of the correct response with an appropriate description for the same. *Here is a summary of the instructions for the main task. In this task, you will be presented with multiple blocks of trials. A fixation cross which looks like this (+) will precede each trial. Please fixate on the cross before the start of the trial. In each trial, you will be presented with two images. Both images may have different levels of contrast (i*.*e. brightness) on each trial. Your task is to choose one of these two images. If you want to choose the image presented on your left, please press the left arrow on your keyboard. If you want to choose the image presented on your right, press the right arrow on your keyboard. Your main aim is to figure out which image you should choose. On each block of trials there is a relationship between the contrast (brightness) level of the image and how often you may win 1 point if you choose that image. For example, on some blocks of trials, the image with higher contrast (brightness) is associated with winning 1 point more often while in another block of trial, the relationship may be reversed. This relationship may change when a new block of trials starts. You will learn this relationship from feedback after your choice. That is, after each trial, you will be presented with the points you win on that trial. You should try to maximize your winnings on each trial. If you have understood the instructions, please press any key to proceed with the experiment*.

An additional set of instructions were presented to participants to explain the use of a slider in Experiment 2.

*Once you make a choice and receive feedback, you will be presented with a slider that ranges between 0 to 100 percent. Again, you will be presented with two images. Both images may have different contrast levels (i*.*e*., *brightness) on each trial. Additionally, one of these two images will have a border. You have to assume that you have hypothetically chosen that image with the border. Based on this hypothetical choice scenario, you are expected to indicate the chance with which you think you can win 1 point on the scale of 0 to 100 percent. Please make the response only when the color of the border changes from red to green. To select the chance, you can drag the slider based on the labels of the slider. You could also directly click on the slider above a particular percent to respond. After your response, please click on the Continue button to proceed in the task. If you have understood these instructions, please press any key to proceed to the experiment*.

This was supplemented with an instructions quiz to ensure thorough understanding of the task, see Instructions quiz. At the end of the task, a debriefing quiz was used to ask participants about the strategies that they used in the task, see Debriefing quiz.

### Instructions quiz for Experiment 1

If you want to choose the image on the left of the fixation (+), which key should you press?

- Right arrow
- Left arrow
- Space Bar

Assume that you won 1 point after choosing the high-contrast image on the left-hand side. Does it mean that you will always win if you choose the image on the left side, irrespective of the contrast levels of the images?

- Yes
- Maybe
- No

Assume that you have previously won 1 point after choosing the image with high contrast in a certain block of trials. Does it mean that you will always win a point when choosing the dark patch in this block?

- No
- May be
- Yes

There could be trials when it can be difficult to distinguish between the patches based on their contrast levels.

- True
- False

Assume that you did not win after choosing the patch that you previously mostly won on. Identify possible reason(s) for it.

- I may have been confused between the contrast levels of the images because they look similar.
- It may happen that I may not win even after choosing the previously rewarding patch because there is no guarantee that you will win on the same patch in a block.
- It is possible that there is no reward associated with both images on certain trials.
- You may have been confused between the contrast levels of the images because they look similar. And it may happen that you do not win even after choosing the previously rewarding patch because there is no guarantee that you will win on the same patch in a block.

Assume that in block 1, the image with higher contrast was almost always rewarding. Does that mean the higher contrast patch will always reward you in the next block?

- Yes
- No

### Instructions quiz for Experiment 2

The percentages on the slider indicate which of the following?

- Chances of winning 0 points, if you chose the image with the green border.
- Chances of winning 1 point, if you chose the image with the red border.
- Chances of winning 2 points, if you chose the image with the green border.
- Chances of winning 1 point, if you chose the image with the green border.

Once you have clicked on the slider to respond, how can you use the slider to re-adjust your response to the desired percent level?

- Directly click on the slider corresponding to the desired percent labels.
- Re-adjustment of response is not possible once the slider has been initially clicked.
- Click on the slider and then drag it to a corresponding desired percent label.

If you think that the chance of winning 1 point is 70 percent when choosing the image with the green border, to which percent label would you drag the slider to?

- 30 percent.
- 60 percent.
- 70 percent.

There may be trials where you win 0 points more often on choosing a certain image and yet, you may be asked to estimate the chances of winning 1 point for the same image using the slider.

- True.
- False.

You are allowed to respond using the slider, when the color of the border is:

- Red.
- Green.
- None of the above.

### Debriefing quiz for Experiment 1

1. 1. Which of these options is correct? (To address the bias)
  - a. I always chose the image with the high contrast level.
  - b. I always chose the image with the low contrast level.
  - c. I chose the low contrast image more often for some blocks of trials, while the reverse was true for other blocks of trials.
2. Assume that you won 1 point by choosing the image on the left side of the fixation (+). Consequently, did you always choose the image on the left side, irrespective of its contrast levels? (Location)
  - a. Yes
  - b. No
  - c. Depended on the block of trials.
3. Assume that you won 1 point in a trial, after choosing a high contrast image. Consequently, did you always choose the image with high contrast in the next trials, in that given block? (Reward Uncertainty)
  - a. Yes
  - b. No
  - c. Depended on the block of trials
4. There were trials in the task in which you were rewarded 0 points, despite choosing the image that you previously won on. (All types of Uncertainty)
  - a. True
  - b. False If true, what do you think were the reasons for the same?
  - a. The images had similar contrast levels and I got confused between them.
  - b. This occurred because there was no guarantee that I would win 1 point even after choosing the image that was previously rewarding.
  - c. The images had no points associated with them.
  - d. There could be multiple reasons for it. I could have been confused because of the similar contrast levels between the images and there is no guarantee that I would 1 point despite choosing the more rewarding image.
5. I found it more difficult to tell the images apart from one another (based on contrast levels), on certain trials. (Perceptual Uncertainty)
  - a. True
  - b. False
6. Assume you won 1 point more often, when you choose the high contrast image in block 1. Did the same happen to you in the next block of trials? (Block)
  - a. Yes, always.
  - b. No, never.
  - c. Sometimes.
7. If you wanted to choose the image on the right side of the fixation (+), which key did you press? (Key Press)
  - a. Right Arrow
  - b. Left Arrow
  - c. Space Bar
  - d. Any other key
8. Imagine that you were responding to a block of difficult trials i.e. when you were not able to figure out the more rewarding image. In such a scenario, did you have a preference to respond towards a particular image? If so, please indicate.
  - a. High Contrast Image
  - b. Low Contrast Image
  - c. No, I had no such preference.

### Debriefing for Experiment 2

I used the slider to indicate the chances of winning 1 point, if I chose the image with the green border.

- True.
- False.
- Sometimes.

